# Cross-regulation between CDK and MAPK control cellular fate

**DOI:** 10.1101/274787

**Authors:** Eric Durandau, Serge Pelet

## Abstract

Commitment to a new cell cycle is controlled by a number of cellular signals. Mitogen-Activated Protein Kinase pathways, which transduce multiple extracellular cues, have been shown to be interconnected with the cell cycle. Using budding yeast as a model system, we have quantified in hundreds of live single cells the interplay between the MAPK regulating the mating response and the Cyclin-Dependent Kinase controlling cell cycle progression. Different patterns of MAPK activity dynamics could be identified by clustering cells based on their CDK activity, denoting the tight relationship between these two cellular signals. In mating mixtures, we have verified that the interplay between CDK and MAPK activities allows cells to select their fate, preventing them from being blocked in an undesirable cellular program.

## Introduction

Cells have developed complex signal transduction pathways to respond to changes in their environment. Plasma membrane sensors detect extracellular stimuli and relay this information inside the cell via signaling cascades. These protein networks can integrate multiple information to launch the appropriate cellular response. For instance, Mitogen-Activated Protein Kinase (MAPK) pathways are central nodes in the signaling network of eukaryotic cells, because they relay extracellular cues such as growth factors or stresses (Roux and Blenis, 2004; Chen and Thorner, 2007). The metabolic state, the cellular morphology or the cell cycle phase of individual cells can be integrated by the MAPK cascade to finely tune the cellular response (Strickfaden *et al*., 2007; Nagiec and Dohlman, 2012; Clement *et al*., 2013). However, the molecular mechanisms that allow these signal integrations are generally unknown.

The simplified settings offered by *S. cerevisiae* provide an ideal platform to study these complex mechanisms. The budding yeast MAPK network is composed of four main pathways active in haploid cells (Saito, 2010). Multiple instances of signal integration have been documented in this model system: the cross-inhibition between two MAPK pathways (Nagiec and Dohlman, 2012), the limitation of signal transmission in low nutrient conditions (Clement *et al*., 2013; Sharifian *et al*., 2015) and the tight coupling between cell cycle regulation and MAPK activity (Oehlen and Cross, 1994; Peter and Herskowitz, 1994; Wassmann and Ammerer, 1997; Escoté *et al*., 2004; Clotet and Posas, 2007; Strickfaden *et al*., 2007). In this study, we were interested in the interplay between the cell cycle and the mating pathway.

Haploid budding yeasts exist in two mating types: *MAT***a** and *MAT***α**. They produce pheromones (respectively α-factor and α-factor) that can be sensed by a mating partner. Activation of a G-protein-coupled receptor by the pheromone leads to the activation of the MAPKs Fus3 and Kss1, via a three-tier kinase cascade recruited to the plasma membrane by the scaffold protein Ste5 (Bardwell, 2005; Atay and Skotheim, 2017). The two MAPKs initiate a mating program that includes the transcription of hundreds of genes, the arrest of the cell cycle in G1, the formation of a mating projection and which culminates in the fusion of the two partners.

In order to guarantee that each cell that undergoes fusion possesses a single copy of its genome, active Fus3 phosphorylates the Cyclin Kinase Inhibitor (CKI) Far1 which arrests the cells in G1 (Peter *et al*., 1993; Peter and Herskowitz, 1994). In addition, during division, signaling in the mating pathway is dampened by the action of the Cyclin Dependent Kinase (CDK) Cdc28. Cdc28, the only CDK in *S. cerevisiae*, associates with the different cyclins to ensure the proper progression through the cell cycle. Inhibition of the mating pathway is made possible by the association between the CDK and the late-G1 cyclins Cln1 and Cln2. The Cln1/2-Cdc28 complex has been shown to phosphorylate the scaffold protein Ste5, thereby preventing its recruitment to the plasma membrane and thereby preventing the transduction of the signal from the receptor to the MAPK cascade (Strickfaden *et al*., 2007).

Most of the knowledge on the mating-induced cell cycle arrest in G1 and the inhibition of the mating pathway during division has been obtained from population-level measurements, relying on artificial synchronizations of the cell cycle using temperature sensitive mutants, chemical inhibitors or by overexpression of cyclins (McKinney *et al*., 1993; Peter *et al*., 1993; Wassmann and Ammerer, 1997; Strickfaden *et al*., 2007). More recent studies have used single cell measurements to monitor this cross-inhibition, but focused either on Ste5 relocation (Repetto *et al*., 2018) MAPK activity (Durandau *et al*., 2015; Conlon *et al*., 2016) or on CDK activity (Doncic *et al*., 2015).

In this study, we have developed a sensitive assay enabling to quantify in parallel MAPK and CDK activity in non-synchronized live single cells using fluorescent biosensors. By exploiting the natural diversity present in the population, we have been able to cluster cells based on their cell cycle position and monitored their MAPK activity pattern. We could confirm the key role of Far1 for the G1 arrest. However, our data suggest that an additional mechanism working in parallel with the Ste5 phosphorylation is required to limit signaling during S-phase. Furthermore, we highlight the importance of the cross-inhibition between MAPK and CDK for cell-fate decision in the mating process. The interplay between these two activities will determine whether cells induce a mating response or commit to a new cell cycle round.

## Results

### Quantifying MAPK activity

In order to quantify mating MAPK activity, we have developed a Synthetic Kinase Activity Relocation Sensor (SKARS). This probe was engineered by combining a specific MAPK docking site, a phosphorylatable Nuclear Localization Signal (NLS) and a Fluorescent Protein (FP) (Durandau *et al*., 2015). The docking site consists in the first 33 amino acids from the MAP2K Ste7, which confers specificity towards both Fus3 and Kss1 (Reményi *et al*., 2005). Under normal growth conditions, the NLS promotes the enrichment of the sensor in the nucleus. Upon activation of the MAPK, specific residues neighboring the NLS are phosphorylated by the targeted MAPK, leading to a decrease in the import rate of the sensor in the nucleus. The presence of the FP allows monitoring nuclear-to-cytoplasmic partitioning of the sensor as function of time (Figure 1A). We use two bright field images and a fluorescence image of the nucleus (Hta2-tdiRFP) to segment the nuclear and cytoplasmic areas in each cell and measure the fluorescence intensity in the other fluorescent images acquired (Pelet *et al*., 2012). The ratio of mean fluorescence intensities between these two compartments in the RFP channel provides a dynamic measure of MAPK signaling activity in each single cell.

**Figure 1:**
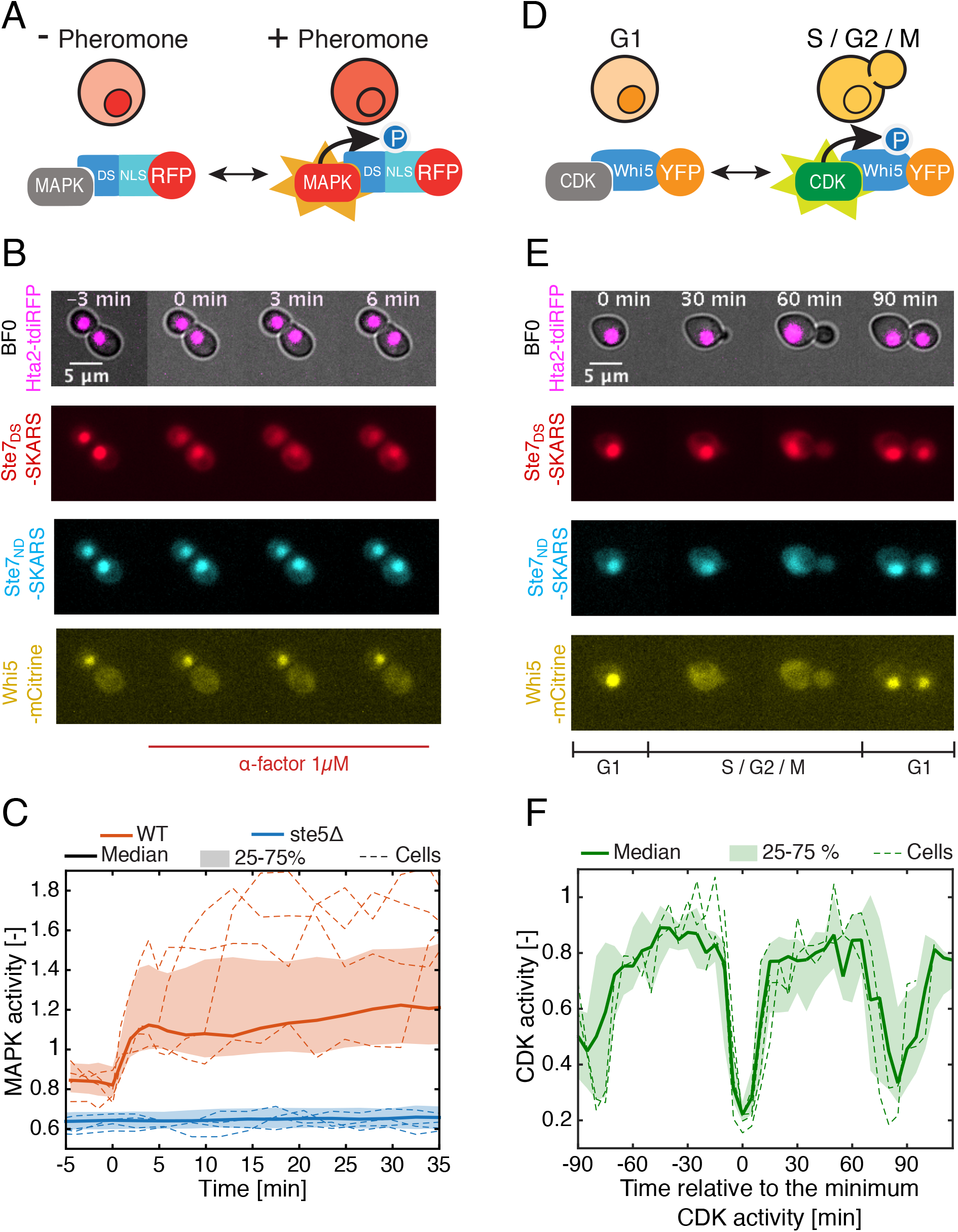
Dynamic MAPK and CDK activity measurements in live single cells. **A.** Schematic representation of the Synthetic Kinase Activity Relocation Sensor (SKARS). **B.** Epi-fluorescence microscopy images of WT cells expressing a Hta2-tdiRFP (nucleus), a Ste7_DS_-SKARS-mCherry (MAPK activity sensor), a Ste7_ND_-SKARS-CFP (Corrector), and a Whi5-mCitrine (CDK activity). Log-phase cells were placed in a microscopy well-plate. During the time lapse, cells were stimulated with 1 μM α-factor (time 0). **C.** MAPK activity was quantified from time-lapse movies acquired with a WT strain (red, Nc = 191) and a mating signaling dead mutant (*ste5* Δ, blue, Nc = 437) (See Methods and Supplementary Figure 1). For all similar graphs, the solid line represents the median of the population, the shaded area delimits the 25 and 75 percentiles of the population. Dashed lines display the response of a few single cells. Nc represents the number of single-cell traces analyzed. **D.** Schematic representation of the CDK activity sensor (Whi5-mCitrine). **E.** Microscopy images of the same strain presented in B. The thumbnails are extracted from a 120 min time-lapse movie of unstimulated cells dividing under normal growth conditions. **F.** CDK activity reported as function of time relative to G1. Single cell traces that exhibit at least one trough of CDK activity were selected (Nc = 70). The traces were synchronized relative to the time at which the CDK activity reaches its minimum value (see Supplementary Figure 2).

One experimental difficulty associated with this sensing strategy is the fact that each cell has a distinct inherent capacity to import the sensor in the nucleus. Therefore, the read-out provided for each cell includes an additional undesired component. In order to correct for this experimental variability, we introduced, in parallel to the functional sensor present in the RFP channel, a non-functional sensor in the CFP channel based on a version of the reporter that cannot bind the kinase (non-docking sensor, Ste7_ND_-SKARS, Figure 1B). We define the MAPK activity as the ratio of cytoplasmic-to-nuclear fluorescence of the functional sensor over the ratio of cytoplasmic-to-nuclear fluorescence of the non-docking corrector (Supplementary Figure 1 and Methods). This metric provides a relative measure of the combined Fus3 and Kss1 activity independently of the nuclear import capacity of each cell.

We have performed time-lapse experiments where cells are stimulated with synthetic pheromone at saturating concentration (1000nM α-factor) directly under the microscope. The dynamics of activation of the mating pathway in single cells can be quantified (Figure 1C). In comparison with a signaling dead mutant (*ste5*Δ), we observe a clear increase in the median MAPK activity of the population within 5 minutes after the stimulus. However, the few single cell traces plotted in this figure display strikingly different dynamic behaviors, demonstrating well the great heterogeneity in the signaling capacity of individual cells.

### Monitoring CDK activity

Because it has been established for a long time that the cell cycle can influence mating signaling competence, we decided to monitor in the same cells both MAPK and CDK activities to overcome the need for population average measurement and artificial cell cycle synchronization. In order to follow cell cycle progression in an automated and robust manner, we used a fluorescently tagged Whi5, which has been used in many studies as an endogenous relocation probe for the G1 state (Bean *et al*., 2006; Doncic *et al*., 2011). Whi5 is enriched in the nucleus of the cells in G1 to repress the expression of specific cell cycle genes. Phosphorylation by the Cln3-CDK complex relieves this repression by promoting the nuclear export of Whi5 (Costanzo *et al*., 2004; de Bruin *et al*., 2004) (Figure 1D and E). Whi5 will only shuttle back into the nucleus as cells re-enter in G1. Thus, in addition to the SKARS and its corrector, we tagged Whi5 with mCitrine and measured the cytoplasmic to nuclear fluorescence intensity in the YFP channel as a proxy for CDK activity (Figure 1F). Note that during the division process, fluctuation in CDK activity levels cannot be quantified using this approach, but the Whi5 probe allows us to precisely monitor the exit from G1 and the entry into G1. In the dataset of a two-hour time-lapse movie, we collected single cell traces that displayed a transient nuclear accumulation of Whi5. The rapid accumulation of Whi5 in the nucleus, which corresponds to the sharp decrease in CDK activity occurring at the onset of G1, was used to synchronize computationally the single cell traces (Figure 1F and Supplementary Figure 2). The alignment of the single-cell responses reveals two additional CDK activity drops taking place roughly 90 minutes before or after the central trough. This timing matches the expected cell cycle length in these conditions (Charvin *et al*., 2008). To sum up, this reporter strain allows us to monitor in parallel CDK and mating MAPK activities in single cells, thereby providing an assay to disentangle their interactions.

### Cell cycle stage clustering

In a naturally cycling population of budding yeast cells, all cell cycle stages are represented at a given time point. Therefore, when this population is stimulated with α-factor, in a single experiment, we can observe the entire diversity of responses present in the population generated by this extrinsic variability. In a typical time-lapse experiment of 45 minutes, cells are imaged every 3 min and the pheromone is added before the third time point. From such a microscopy dataset, several hundreds of single cells are tracked and pass through a quality control filter. Note that unless stated otherwise, all following experiments with exogenous pheromone stimulation are performed in a *bar1*Δ background, to maintain a constant extracellular concentration of α-factor in the medium surrounding the cells and were performed in at least three independent experiments.

The single cell traces are clustered *in silico* based on their CDK activity pattern to identify the different cell-cycle stage populations (see Methods). The first cluster consists of cells with low CDK activity throughout the time lapse (*G1*, 20% of the population). These *G1* cells are MAPK signaling competent and display a strong and sustained response of the SKARS upon addition of pheromone (Figure 2A). A second cluster is made of cells with a high CDK activity during the 45 minutes of the time lapse (*Out-of-G1*, 35%). These cells display a fast response to the mating pheromone but with a low amplitude. Previous reports have shown, by overexpression of cyclins, that the activity of the CDK can abolish or strongly reduce the activity of the mating pathway (Oehlen and Cross, 1994; Wassmann and Ammerer, 1997). Under physiological conditions, our data show that dividing cells are none-the-less able to activate the MAPK pathway, but only weakly compared to the *G1* cells.

**Figure 2:**
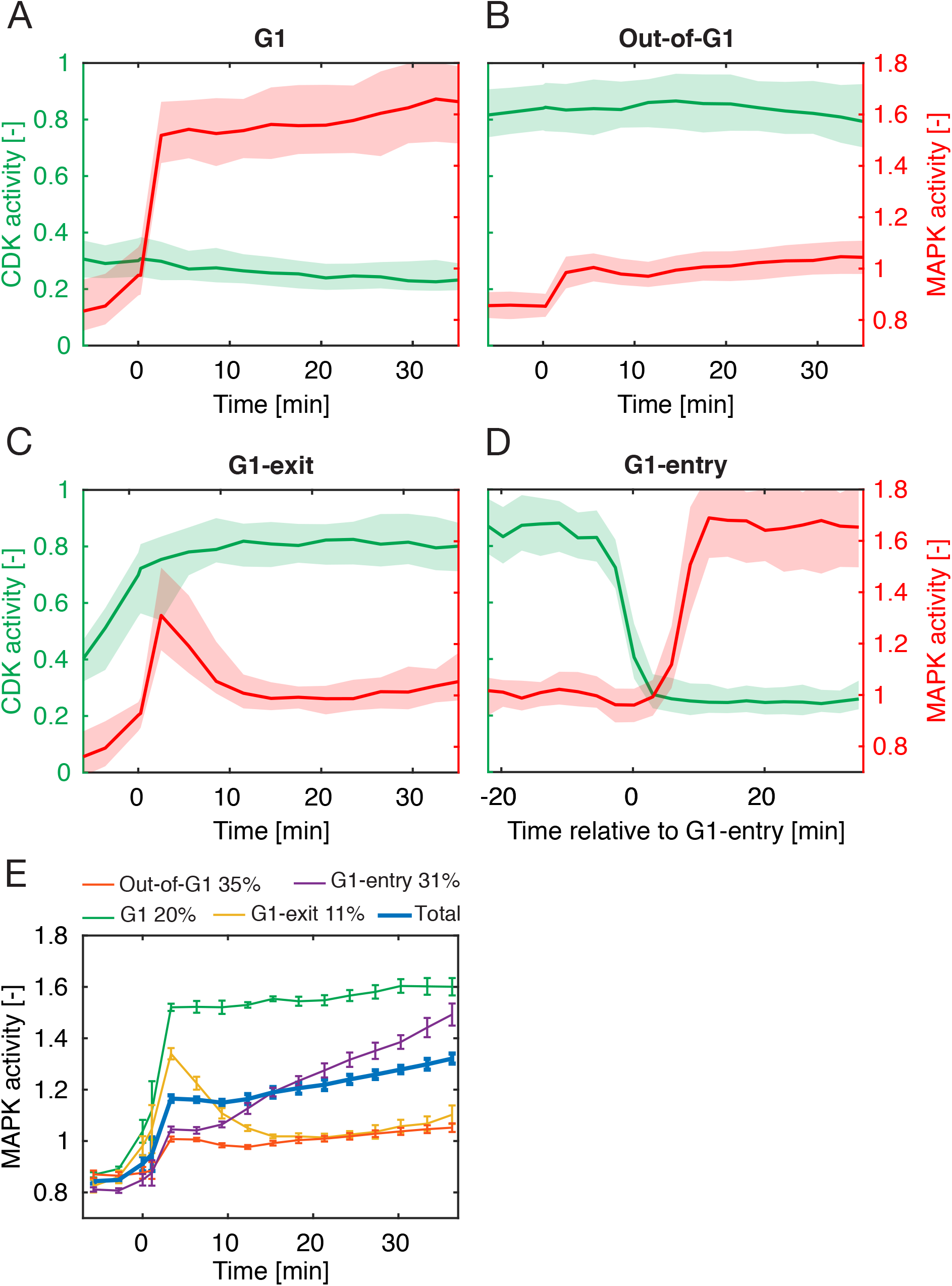
Dynamics of MAPK activity monitored by the SKARS in different cell cycle phases. **A-D.** MAPK activity was quantified for cells in an asynchronously growing culture stimulated with 1 μM α-factor at time 0. 738 single cell traces were clustered by comparing the CDK activity prior and after stimulation (see Methods). *G1* cells (A, Nc =143) retain a low CDK activity throughout the time lapse, while *Out-of-G1* cells (B, Nc =256) maintain a high CDK activity. *G1-exit* cells (C, Nc =76) start with low and finish with high CDK activity. Conversely, *G1-entry* cells (D, Nc =249) start with high and finish with low CDK activity. In this sub-population, the traces are aligned relative to the time of G1-entry. The median (solid line) MAPK activity (red) and CDK activity (green) and the 25–75 percentiles (shaded area) of each sub-population are plotted. **E.** Summary of the various dynamic MAPK activities observed upon addition of 1 μM α-factor at time 0 in the four main cell cycle clusters. Each line corresponds to the mean of the medians of 5 biological replicates. Error bars represent the standard deviations of the medians. The percentages in the legend indicate the relative proportion of each cluster in the population.

In addition, our dynamic measurements can reveal interesting signaling patterns by focusing our analysis on cells that transition between these two strong and weak signaling states (*G1* and *Out-of-G1*, respectively). A third cluster was thus defined with cells that start with a low CDK activity and end with high CDK activity. These cells are thus exiting G1 and enter S-phase at some point during the time lapse (*G1-exit* 11%, Figure 2C). Interestingly, these cells respond strongly to the pheromone stimulus, but display a transient MAPK activation behavior. When pheromone is added, the mating pathway is first rapidly activated. However, when the CDK activity builds up in the cell, an inhibition of the MAPK pathway sets in, leading to a return of the MAPK activity to a low level roughly 15 min after the stimulus.

The fourth cluster consists in cells that enter in G1 during the time lapse (*G1-entry*, 31%). These cells start with high CDK activity and end the time lapse with low CDK activity. This transition can happen at any time point during the experiment. The median MAPK activity in this sub-population increases gradually with time, while the individual traces display sharp transitions from a low signaling to a high-signaling state (Supplementary Figure 3A). Using the Whi5 reporter, the single cell traces were temporally aligned based on the time of G1 entry. As a result, a synchronous increase in MAPK activity is observed when CDK activity stabilizes to low values, corresponding to the G1-phase (Figure 2D and Supplementary Figure 3B). Finally, a fifth cluster was identified with cells that cycle briefly in G1 during the time lapse (*Through-G1*, 3%). No specific behavior in MAPK activity was observed in these cells even when traces are aligned relative to the time of G1 entry (Supplementary Figure 4).

Figure 2E allows a direct comparison of the MAPK activity measured with the SKARS in different cell cycle stages. The mean response of the population (thick blue line) is not representative of the behavior of individual cells which can display strikingly different behavior. However, because CDK plays a major role in controlling the signaling output of the mating pathway, clustering based on the CDK activity pattern allows to group together cells that display similar MAPK activity dynamics. Thus, we observe four types of behaviors: strong and sustained activity for *G1* cells, weak and sustained activity in *Out-of-G1* cells, strong but transient activity in the *G1-exit* cluster and delayed activation (depending on the timing of CDK activity drop) present in the *G1-entry* cells.

### Comparison with other reporters

In order to compare the results obtained with the SKARS, we performed a similar analysis with two different assays that report on mating pathway activity. Using a fluorescently tagged Kss1 (Kss1-mScarlett), the relocation of the MAPK from the nucleus to the cytoplasm upon pheromone stimulus has been monitored in a strain carrying Whi5-mCitrine. This change in cellular compartments of Kss1 has been shown to be a consequence of the disassembly of a complex formed between Dig1/Dig2, Ste12 and Kss1, upon phosphorylation by Fus3 and/or Kss1 (Pelet, 2017) (Supplementary Figure 5 A and B). When comparing the response of the cells in different cell-cycle stages, a similar pattern can be observed between this assay and the SKARS reporter (Supplementary Figure 6). The major difference can be observed in the *G1-entry* cluster where a gradual increase in signaling activity can be seen upon entry into G1 (Supplementary Figure 6D), compared to the sharper dynamics of the SKARS reporter in the same cell cycle stage (Figure 2D). In addition, when comparing the Kss1 relocation of *G1* and *G1-entry* cells relative to the time of pheromone addition (Supplementary Figure 6E), we see that they are almost identical. This suggests that cells late in the division process have fully recovered their signaling ability.

The second assay is based on the dPSTR (dynamic Protein Synthesis Translocation Reporter) system which allows to monitor the dynamics of induction of a promoter of interest (Aymoz *et al*., 2016). This promoter drives the expression of a small peptide that promotes the relocation of a fluorescent protein in the nucleus of the cell. We use the previously published pAGA1-dPSTR to monitor the dynamics of mating gene induction. AGA1 has been shown to be strongly induced by α-factor in a MAPK dependent manner (Roy *et al*., 1991; Oehlen *et al*., 1996; Aymoz *et al*., 2018). Clustering of the gene expression data based on cell-cycle stage demonstrates a strong expression in *G1* and *G1-entry* clusters, while clusters for *Out-of-G1* and *G1-exit* cells display an attenuated and delayed response (Supplementary Figure 7).

The global outcome that can be obtained by comparing these three types of reporters is that full signal competence is observed in the *G1* cluster, while *Out-of-G1* cells have a reduced ability to signal. In addition, cells exiting G1 will only transiently activate the MAPK. Interestingly, this transient activation is not sufficient to drive gene expression as protein production in the *G1-exit* cluster is delayed by 20 to 30 minutes.

### MAPK activity in the G1-entry cluster

The comparison between the SKARS, the Kss1 relocation and the pAGA1-dPSTR also highlights a discrepancy for the *G1-entry* cluster. While the two latter assays display a signaling ability that is comparable to the one present in *G1* cells for the *G1-entry* cluster, the SKARS measurements suggest that the MAPK activity is only recovered when CDK activity has dropped, as cells enter in G1.

The influence of the cell cycle on the nuclear enrichment of the SKARS could potentially explain this observation. Indeed, a lower enrichment of the corrector can be observed in *Out-of-G1* cells compared to *G1* cells (Supplementary Figure 8A and B). In the *G1-entry* cluster, this results in a slow transition from a low to a high nuclear to cytoplasmic ratio (N/C) of the corrector around the time of entry into G1. In signaling dead cells, the exact same behavior is observed for the functional sensor (Supplementary Figure 1B). However, the dynamic of this transition is strikingly different from the dynamic of MAPK activity measured upon *G1-entry*. While the corrector slowly rises from −10 to +10 min, the MAPK activity shift takes place between +5 and +10 min after the CDK activity drop. Thus, the fact that these two events are not synchronous, strongly suggests that the sharp increase in MAPK activity is not an artifact from a reduced sensitivity during the division of the cells.

Because, in each individual cell, the sensor and corrector nuclear enrichment are highly correlated (Supplementary Figure 8C), we performed an additional verification and separately analyzed cells with low and high nuclear enrichment of the corrector (Supplementary Figure 8D and E). If the G1-entry behavior was an artifact due to a poor enrichment of the sensor, we would expect that cells with a relatively high nuclear enrichment would not display this behavior or at least to a lower extent. On the contrary, we observe that *G1-entry* cells which keep a high N/C of the corrector throughout the time-lapse display a stronger change in MAPK activity 10 minutes after G1 entry.

In agreement with the Kss1 and dPSTR assays, previous works (Oehlen and Cross, 1994; Wassmann and Ammerer, 1997; Strickfaden *et al*., 2007; Conlon *et al*., 2016), have shown that CDK inhibition of the mating pathway is limited to the S-phase and full signaling competence is recovered in G2-M. The dynamics observed with the SKARS are therefore unexpected. This behavior could be explained by the action of phosphatases acting on the phosphorylated residues present on the SKARS. For instance, the Cdc14 phosphatase leaves the nucleus during mitotic exit to dephosphorylate cyclin-dependent targets (Shou *et al*., 1999; Visintin *et al*., 1999; Mohl *et al*., 2009). If this hypothesis is true, we can envision that other substrates of the MAPK might be the target of phosphatases upon G1-entry and restrict their activity to the G1 phase.

### Pheromone dose response

In order to test how the MAPK signaling behavior changes as function of input strength, time-lapse movies were recorded with various concentrations of pheromone. The same cell cycle clustering approach was used for all the dataset. As expected, the amplitude of the MAPK activity decreases with lower pheromone concentrations (Figure 3 A-C). However, other changes can also be observed between the different concentrations. In the G1-phase, a sustained MAPK activity is observed at high doses of pheromone, while the signaling activity declines at lower α-factor doses (Figure 3A). This behavior suggests an interplay between positive and negative feedback loops. Multiple regulatory mechanisms have been identified in the mating pathway (Hao *et al*., 2003; Bhattacharyya *et al*., 2006; Yu *et al*., 2008; Nagiec *et al*., 2015). None-the-less, it remains difficult to estimate their relative influence on the signaling outcome. Our data suggest that at low α-factor concentrations, negative feedbacks are prevalent and contribute to the deactivation of the pathway. At high pheromone concentrations, however, the positive feedbacks stabilize the system in a high activity state for a long time. Similar experiments were performed in *BAR1*+ cells, where the presence of the pheromone protease adds another layer of regulation to the system and leads to a faster decline in signaling activity at all concentrations but the highest one, which remains sustained over the course of the time lapse experiment (Supplementary Figure 9).

**Figure 3:**
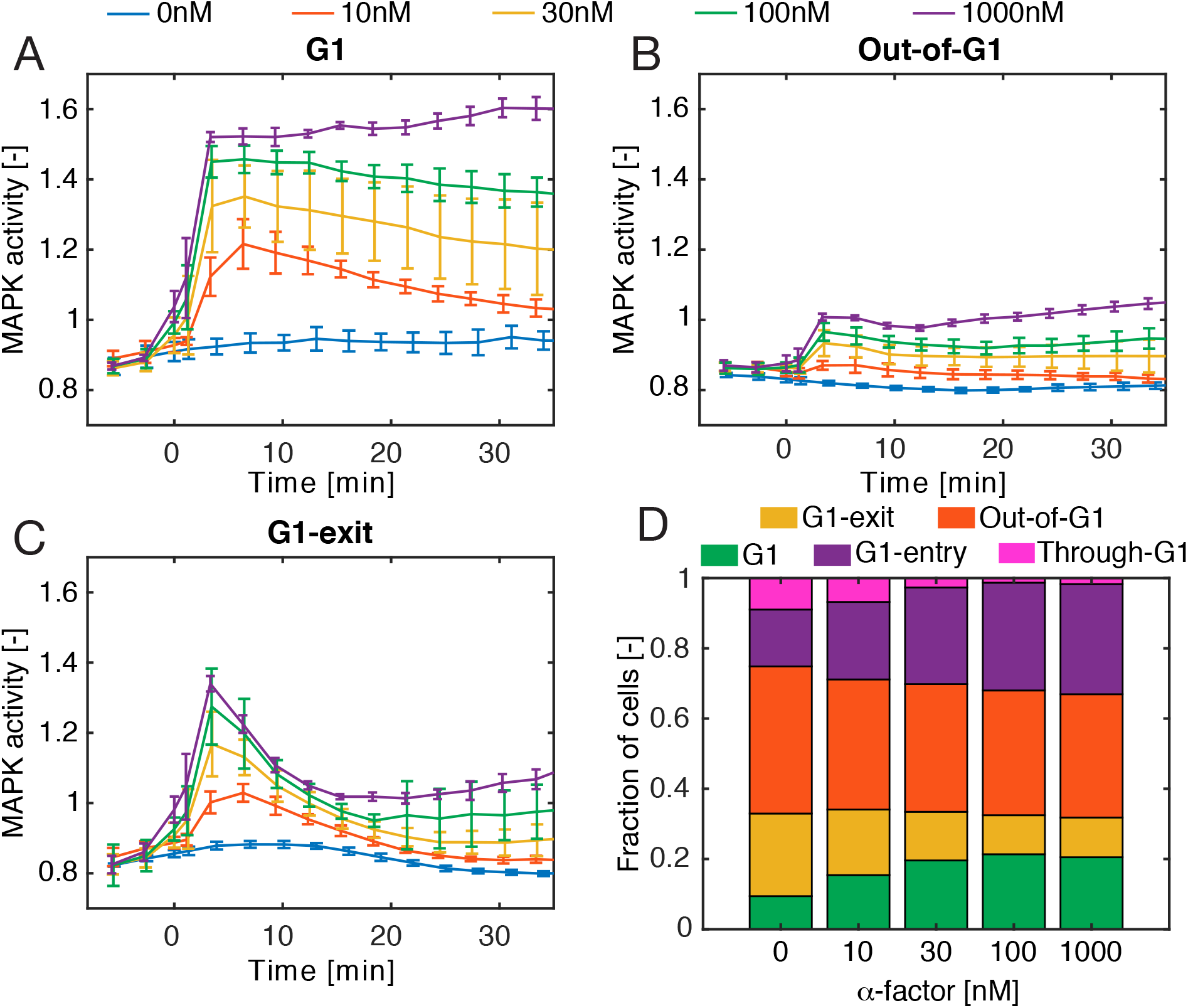
Commitment to cell cycle is pheromone dose-dependent. **A-C.** Pheromone dose-dependent dynamic MAPK activity in different phases of the cell cycle. Experiments and clustering analysis were performed as in Figure 2. Different α-factor concentrations were used to stimulate cells. Median MAPK activity from at least five replicates were averaged for each curve. The error bars represent the standard deviations between the replicates. **D.** Fractions of cells in each sub-population as function of the pheromone concentration for our 45 min time lapse. The prevalence of each cluster is calculated from the means of at least three replicates.

It is well-established that treatment with pheromone prevents cells from entering a new cell cycle round (Hartwell *et al*., 1974), therefore one expects a difference in the proportion of G1 cells at the end of the experiment between α-factor treated and mock treated cells (Figure 3D). Interestingly, we noticed a gradual increase in the fraction of cells retained in G1 as function of pheromone concentration. If we focus on the population of cells starting in G1 (*G1* and *G1-exit* clusters, which roughly represent one third of the population), in the untreated situation 20% will exit G1, while 9% remain in G1 for the entire time lapse. Note that these measured percentages are specific to our 45 min the time lapse. For the sample stimulated with pheromone, this proportion gradually increases to reach 20% of cells that remain in G1 at 1000nM versus 10% that can still escape G1 arrest. A similar behavior is observed for cells that are dividing at the onset of the time lapse (*Out-of-G1* and *G1-entry*). The proportion of the cells that enter and stay in G1 (*G1-entry*) evolves from 12% to 30% as pheromone concentration increases. Thus, the stronger MAPK activity measured at high concentrations promotes a larger accumulation of cells in G1 at the end of the time-lapse movie. These experiments clearly illustrate the fact that the level of MAPK activity influences the ability of the cells to initiate division.

### START as a signal integration point

The START event in the cell cycle has been operationally defined as the time when cells become insensitive to a pheromone stimulus and are committed to a new cell cycle round (Hartwell *et al*., 1974). Multiple cellular events are coordinated around this decision point. Activity of the G1 cyclin Cln3 increases. It triggers the exit of Whi5 out of the nucleus, thereby allowing the transcription of Cln1 and Cln2. These two cyclins will in turn drive the transcription of downstream S-phase genes (Dirick *et al*., 1995; Costanzo *et al*., 2004; de Bruin *et al*., 2004). We have shown that the number of cells that are blocked in G1 or commit to a new cell cycle round is dependent on the pheromone concentration and thus on the MAPK activity. In parallel, we observed that the CDK activity, estimated from the Whi5 nuclear enrichment levels, gradually increases for cells that commit to a new cell cycle round (*G1-exit* cluster, Figure 4 A-D). At low pheromone concentration, cells can often override the mating signal and enter the cell cycle. However, at high concentrations of α-factor only cells that have already reached a sufficient CDK activity will be able to counterbalance the mating signal. In these cells, the CDK promotes the entry in a new division round. Interestingly, at saturating α-factor concentrations, some cells display a transient activation of the CDK, suggesting that the addition of the pheromone inhibited the progression of the cell cycle (Figure 4E). A behavior that has been previously characterized by Doncic et al. (Doncic *et al*., 2011). Taken together these results comfort the idea that START is a signal integration point where the cells balance the relative MAPK and CDK activities to determine their fate: division or mating.

**Figure 4:**
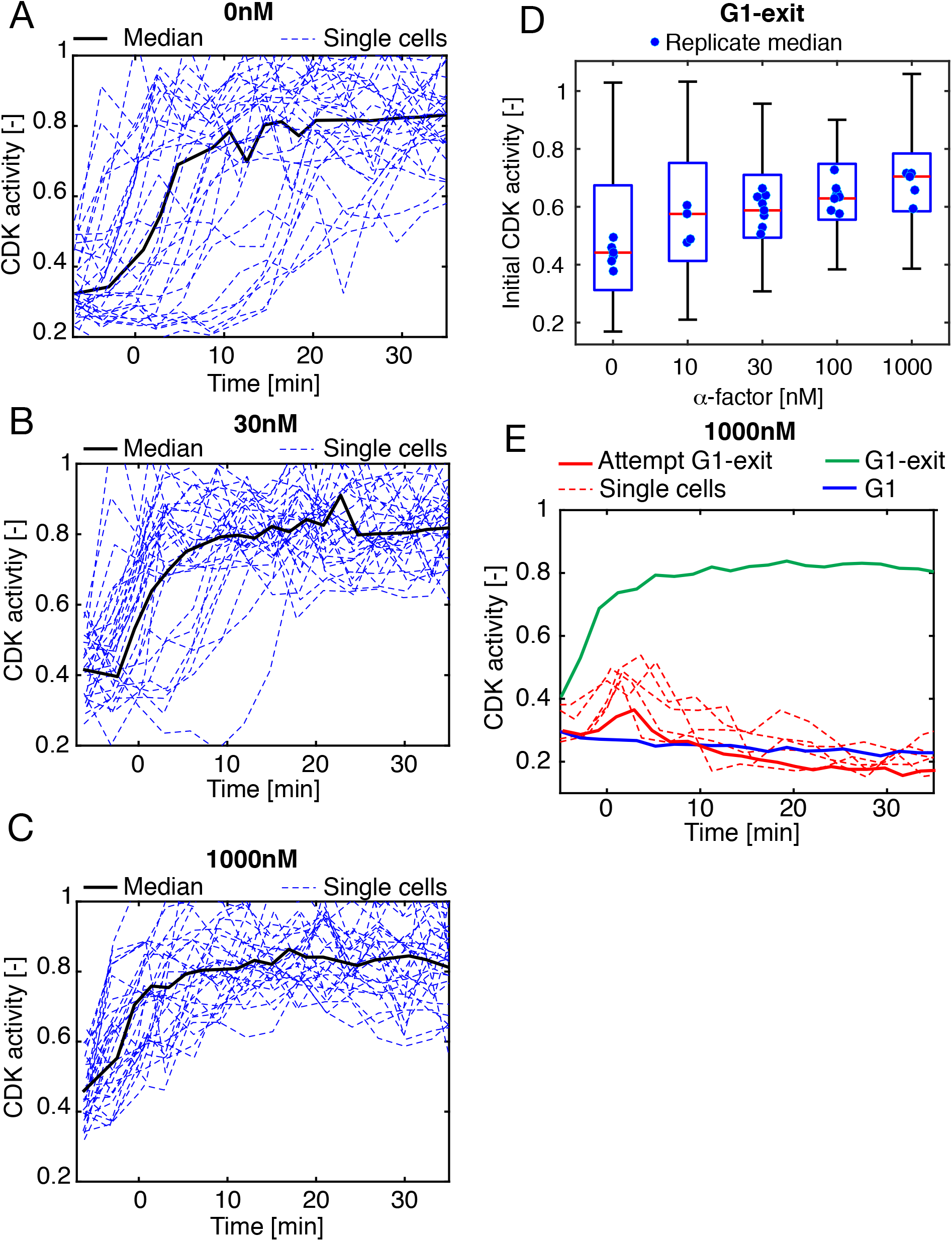
Level of MAPK activity at START controls cell cycle commitment. **A-C.** Dynamic CDK activity of cells exiting G1 at various pheromone concentrations. Single cell traces (dashed lines) and the median of the sub-population (solid line) are presented in each graph. Note the decreased variability in CDK activity patterns as pheromone concentration increases. **D.** Initial CDK activity (mean of the three points before stimulus) measured in the *G1-exit* cluster for different α-factor concentrations. The distribution of the initial CDK of *G1-exit* cells for one replicate for each concentration is presented as a boxplot. The dots correspond to the median initial CDK activity of additional replicates. **E.** CDK activity of cells attempting to exit G1. The dynamic CDK activity of G1 cells from multiple 1000nM α-factor experiments were pooled. The Attempt G1-exit sub-population (red) contains cells with an initial CDK activity above 0.35. Four single cell traces from the Attempt G1-exit sub-population (dashed lines) that display a transient peak in CDK activity around the time of pheromone addition are plotted. The median CDK activity for the *G1* (blue) and *G1-exit* (green) clusters from Figure 2 are plotted for comparison.

Hence, the cross-inhibition between the MAPK and CDK plays a central role in the decision to mate or divide. The molecular mechanisms regulating the interplay between mating and cell cycle have been extensively studied. Phosphorylation and expression of Far1 upon α-factor stimulus inhibit the Cln1/2-Cdc28 complex to prevent cell cycle progression into S-phase (Chang and Herskowitz, 1990; Tyers and Futcher, 1993; Peter and Herskowitz, 1994). Deletion of FAR1 does not affect the MAPK signaling activity of *G1* nor of *Out-of-G1* cells (Supplementary Figure 10A). However, the fraction of cells found in G1 at the end of the time-lapse experiment is increasing from 25% to 50% for WT cells when changing the pheromone concentration to 1μM, while it remains below 30% in *far1*Δ (Supplementary Figure 10B). These results confirm the important role played by Far1 in the G1-arrest during the mating response.

### Modulating Cdc28 activity

We have shown above that the efficiency of the cell cycle arrest depends on the level of MAPK activity. We next want to verify how the CDK activity influence the MAPK signal transduction. In budding yeast, a single Cyclin Dependent Kinase, Cdc28, associates with the different cyclins throughout the cell cycle to orchestrate the division of the cell. While Cdc28 is an essential protein, it is none-the-less possible to acutely inhibit its kinase activity by using an analog sensitive allele (*cdc28-as* (Bishop *et al*., 2000)). The inhibitor NAPP1 was added to the cells 6 minutes after the pheromone stimulus. Upon NAPP1 treatment, *Out-of-G1* cells show a rapid relocation of Whi5 in the nucleus, attesting the fact that CDK activity is blocked. In parallel, the MAPK activity, that is low during division, increases to a level comparable to the one observed in cells stimulated in G1 (Figure 5A).

**Figure 5:**
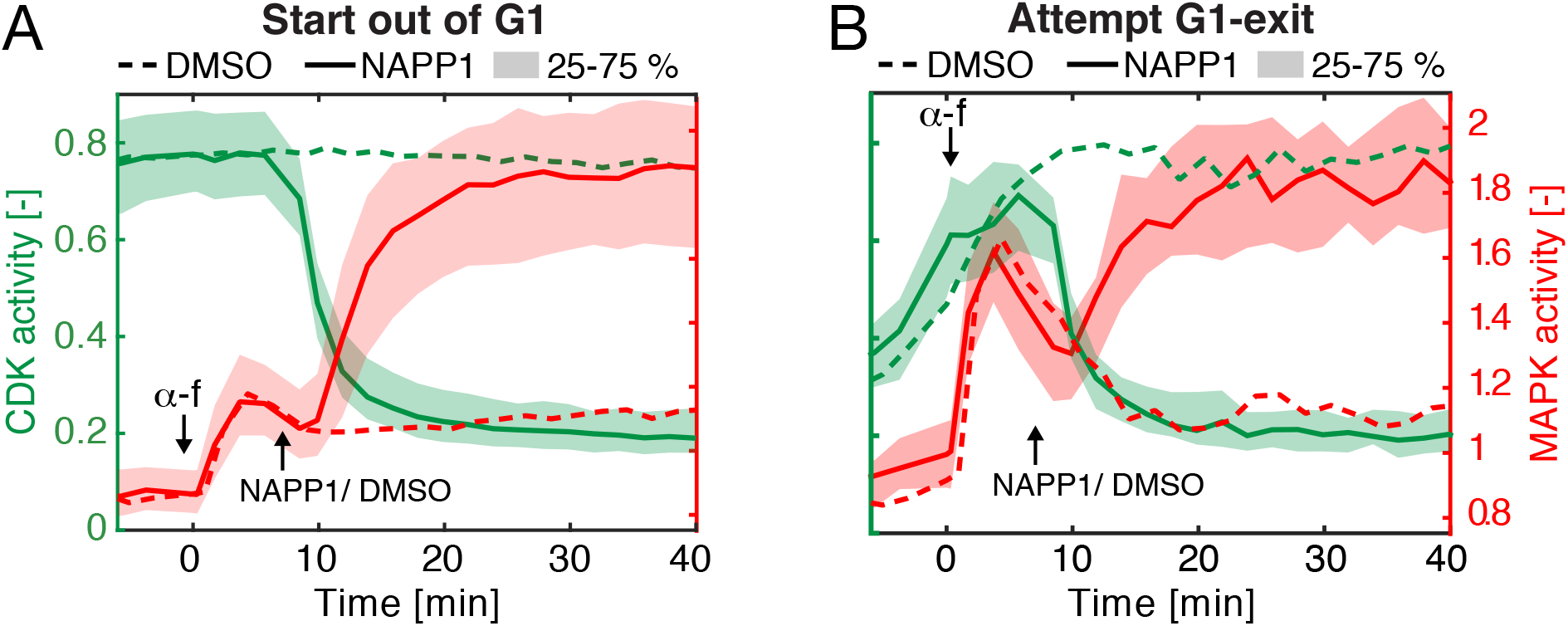
Regulation of MAPK signal transduction by the CDK. **A-B.** Dynamic CDK and MAPK activities after *cdc28-as* chemical inhibition. Cells expressing an analog sensitive allele of Cdc28 (*cdc28-as*) are imaged every two minutes for 50 minutes. Cells are stimulated with 100nM α-factor at time 0. Six minutes later either NAPP1 (10 μM, solid line and shaded area) or DMSO (0.04% v/v, dashed line) is added. MAPK and CDK activities of cells starting *Out-of-G1* (mean CDK_t <0_ >0.5, A, NAPP1: Nc = 448, DMSO: Nc = 381), or exiting G1 upon pheromone addition (mean CDK_t <0_ <0.5 and mean CDK_0 <t <6_ >0.5, B, NAPPI: Nc = 39, DMSO: Nc = 16) are plotted.

Additionally, this experiment allows to verify that the transient MAPK activation observed in the *G1-exit* cluster is shaped by the rising CDK activity. Cells with an increasing CDK activity after pheromone stimulus were clustered. In the DMSO control experiment, this sub-population displays the expected fast activation of the MAPK upon α-factor addition followed by a decay to basal activity as CDK activity rises. In the inhibitor treated cells, the MAPK activity decay is blocked upon NAPP1 addition and the MAPK signal rises to reach full activity (Figure 5B). These experiments demonstrate that the activity of Cdc28 directly and quickly regulates the level of MAPK activity present in the cell.

### Ste5 inhibition by the CDK

One mechanism of inhibition of the mating pathway by the CDK has been shown to consist in the direct phosphorylation of the scaffold Ste5 by the Cln1/2-Cdc28 complex (Strickfaden *et al*., 2007). Eight consensus phosphorylation sites in the vicinity of a plasma membrane binding domain (PM) in Ste5 are targeted by the CDK to alter the charge of this peptide. This phosphorylation prevents the association of Ste5 to the membrane, thereby blocking the signal flow in the pathway. In *ste5*Δ cells, we reintroduced three different alleles of Ste5 at the endogenous locus: the WT copy, the Ste58A mutant (where all putative phosphorylatable residues were mutated to alanine (Strickfaden *et al*., 2007)) or a Ste5_CND_ (where the docking motif of Cln1/2 on Ste5 has been mutated (Bhaduri and Pryciak, 2011)) (Figure 6A). We monitored the response of the mating pathway for these three alleles with the SKARS, the relocation of Kss1 and the p*AGA1*-dPSTR at 10 and 100 nM α-factor (Figure 6B, C and D). The wild-type *STE5* behaved similarly to the WT parental strains, demonstrating the functionality of the complementation. As expected, signaling activity in *G1* cells is minimally influenced by either mutation in Ste5 because the CDK is inactive. In the *G1-exit* cluster, the inhibition following the transient activation of the pathway is less pronounced in the two Ste5 mutants than for the wild-type allele. Generally, the behavior is more pronounced at 10nM than at 100nM α-factor and the Ste5_8A_ mutant displays a weaker inhibition by the CDK than the Ste5_CND_ mutant.

**Figure 6:**
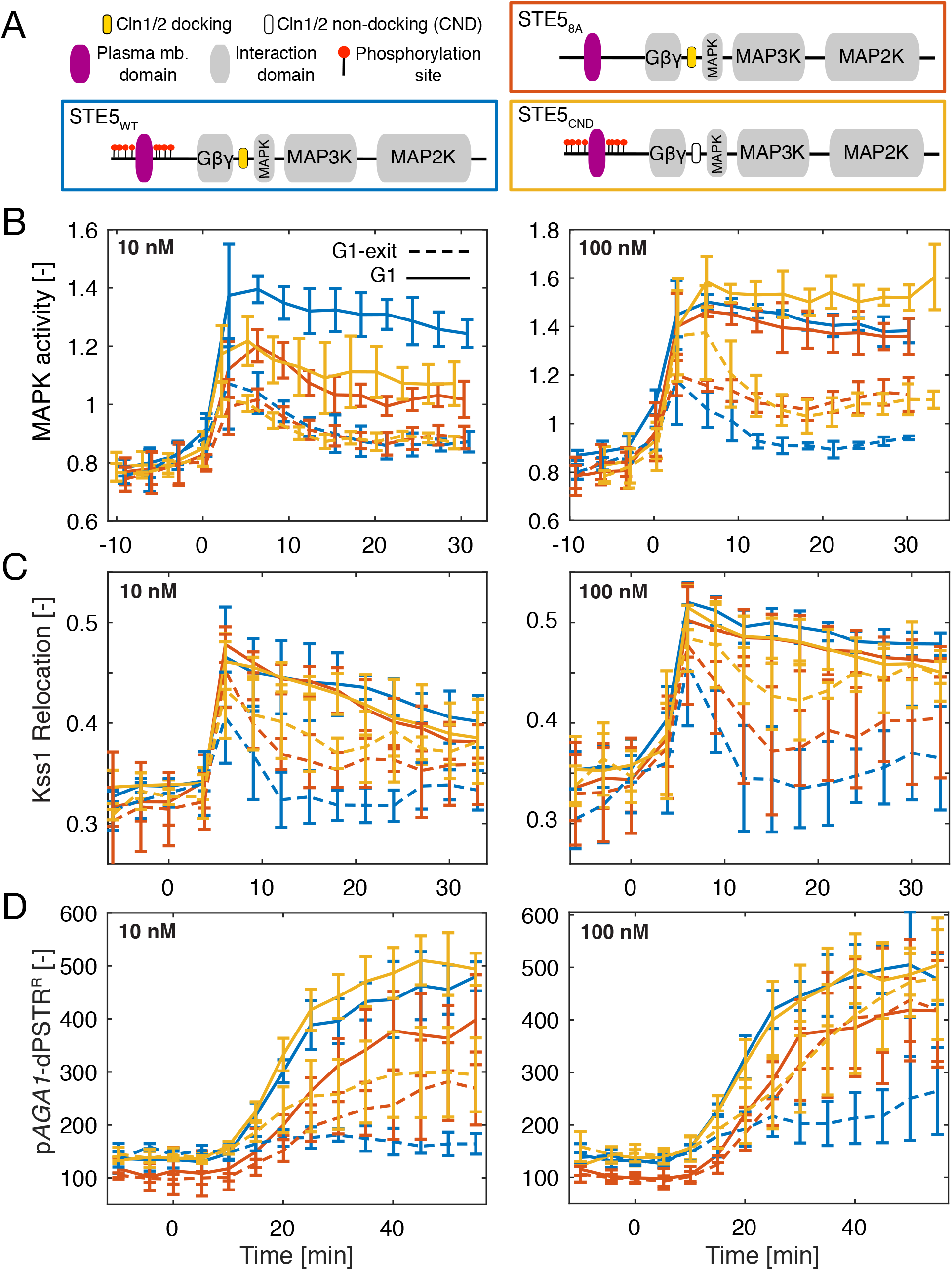
Dynamic MAPK activity of cells expressing non-phosphorylatable Ste5 alleles. **A.** Schematic representation of the wild-type Ste5 scaffold protein structure compared to the Ste5_8A_ mutant with 8 non-phosphorylatable residues in the vicinity of the plasma membrane domain and the Ste5_CND_ variant with point mutations in the Cln1/2 docking motif. **B-D.** Dynamics of mating pathway activity in *G1* (solid line) and *G1-exit* cells (dashed line) measured by the SKARS (B), Kss1 relocation (cytoplasmic over nuclear intensity, C) and p*AGAl*-dPSTR^R^ nuclear accumulation (nucleus minus cytoplasmic intensity, D). Cells expressing either the Ste5_WT_ (blue), Ste5_8A_ (red), or Ste5_CND_ allele (yellow) were treated with 10nM (left) and 100nM (right) pheromone. Cell cycle clustering was performed as in Figure 2. Median MAPK activity from at least three replicates were averaged for each curve. The error bars represent the standard deviations between the replicates.

A recent report has demonstrated that the combined action of Fus3 and Cdc28 is required to phosphorylate the eight residues Ste5 in the vicinity of the PM domain (Repetto *et al*., 2018). However, our data suggest that an additional mechanism might contribute to limit the signal transduction of the mating pathway in the early stage of the division process. This phenomenon is best observed at low concentrations of pheromone where the activity of the MAPK cascade is weaker. This behavior has not been detected previously, probably because most experiments have been performed at saturating levels of pheromone. However, in mating mixture where pheromone concentration is low, this mechanism could contribute to the cell fate decision. In order to identify other potential targets of the CDK, we have tested various alleles of Ste20 because it has often been suggested as a potential target for this regulation (Oehlen and Cross, 1998; Wu *et al*., 1998; Oda *et al*., 1999) (Supplementary Figure 11A and C) and we have also tested the influence of phosphorylation sites on Ste7 without measuring any detectable changes in signaling activity between mutants (Supplementary Figure 11B and D).

To summarize, START is the integration point where MAPK and CDK activities are compared to engage in a specific cellular fate. Far1 and Cdc28 are key players in this cross-inhibition of the two pathways. *far1*Δ cells cannot arrest their cell cycle at START. None-the less, a number of Far1-independent mechanisms for cell cycle arrest have been documented (Tyers, 1996; Oehlen *et al*., 1998; Cherkasova *et al*., 1999). Cdc28 is solely responsible for the inhibition of the mating pathway by the cell cycle and, up to now, only a single target, Ste5, has been convincingly shown to regulate this process. Our data indicate that other mechanisms, that remain to be identified, contribute to the repression of MAPK activity in dividing cells.

### MAPK activity in mating

After studying the response of cells stimulated by exogenous pheromone, we next wanted to understand how the cell cycle and the mating pathway were regulated during mating and how this cross-inhibition allowed an efficient cell-fate selection in these physiological conditions. In order to achieve this, we have imaged the *MAT***a** cells bearing the SKARS, the corrector and the Whi5 marker in presence of a *MAT***α** partner expressing constitutively a cytosolic CFP at high levels. In Figure 7A, thumbnails of such an experiment are displayed. The fusion events (arrow heads) can be detected by observing a sudden increase in CFP signal in the *MAT***a** cells. In the frame preceding the two fusion events displayed in Figure 7A, we observe that the *MAT***a** cells of interest are in G1 (nuclear Whi5) and display a high MAPK activity (nuclear depletion of the SKARS).

**Figure 7:**
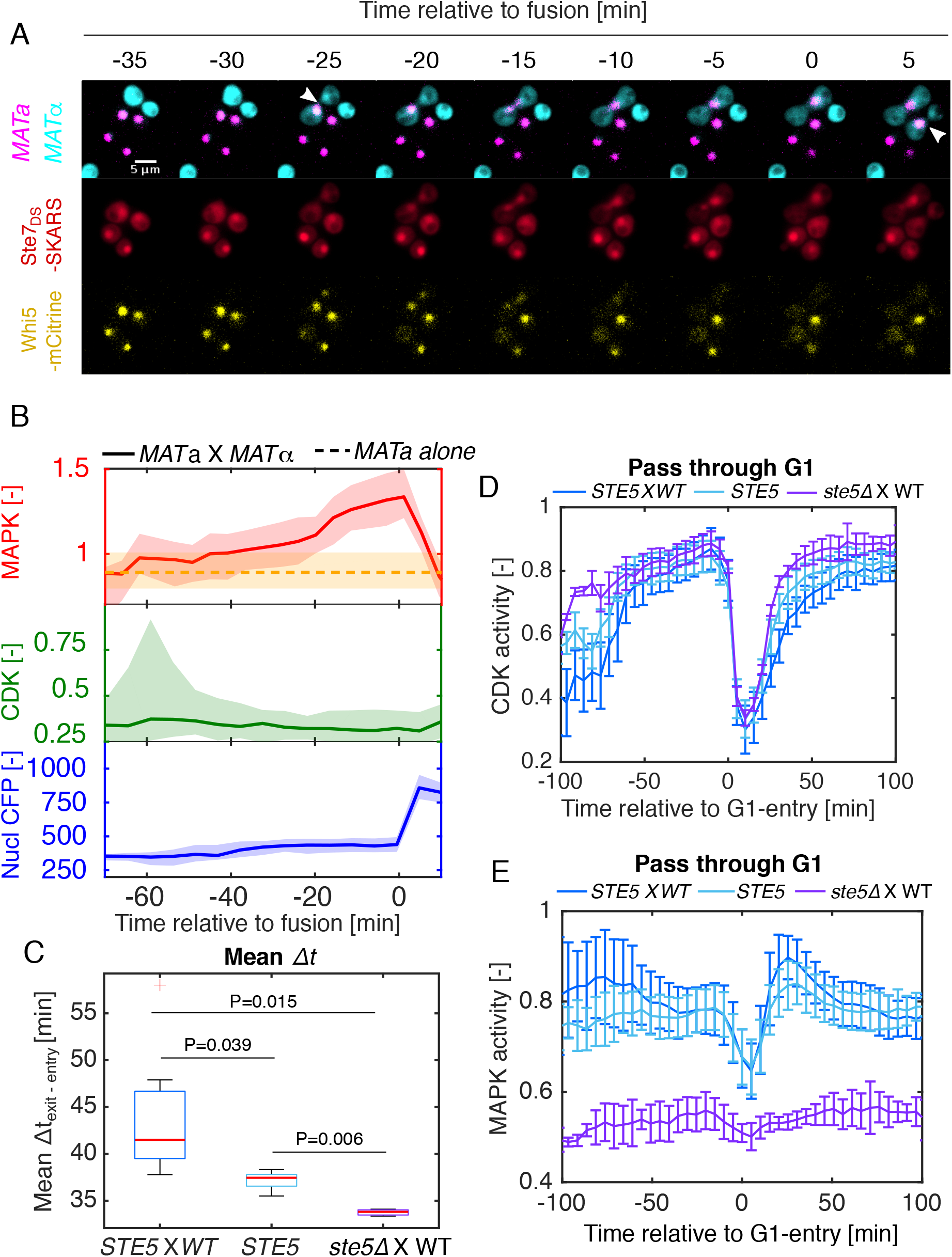
CDK and MAPK cross-regulation in mating conditions. **A.** Microscopy time-lapse images of two fusion events between *MAT***a** cells expressing the various sensors mixed with *MAT***α** strain expressing a cytoplasmic CFP. The fusion can be detected when the CFP from the *MAT***α** diffuses in the *MAT***a** partner (White arrows). The indicated time is relative to the last time point preceding the second fusion event. **B.** Dynamic MAPK and CDK activities in *MAT***a** cells undergoing fusion detected by a sharp increase in CFP intensity (blue) (See Method). MAPK (red) and CDK (green) activities of the fusing cells were synchronized relative to the last time point before the fusion occurs (Nc=127). The dashed orange line and the shaded area around it correspond to the median and 25–75 percentiles (averaged across multiple time points) of the MAPK activity in *MAT***a** cells in absence of mating partners, but imaged in the same conditions. **C-D.** Duration of the G1 phase of non-fusing cells. Non-fusing cells transiting through G1 were identified as previously (Figure 3C and Methods). **C.** The time separating the G1 entry and G1 exit is calculated from each single cell trace (Δt_exit – entry_) and averaged. The means Δt_exit – entry_ from at least three replicates are plotted as boxplot. **D.** The dynamic CDK activity from single cells are synchronized relative to the G1 entry. Error bars are standard deviation of at least three medians replicates. **E.** MAPK activity in non-fusing *MAT***a** cells transiting through G1. Single cell traces are synchronized relative to the G1 entry and averaged as in Figure 2D. Error bars are standard deviation of at least three medians replicates. In these mating experiments, all strains are BAR1.

The dynamic measurements of the mating process allowed us to monitor how cells reach this state. Using our automated image analysis pipeline, we have detected more than one hundred fusion events and curated them manually to remove any artifacts due to a mis-segmentation of the cells. The single cell traces of these events have been computationally aligned relative to the fusion time. Time zero corresponds to the last frame before the increase in CFP intensity, because it is the last time point where MAPK activity can be reliably quantified (Figure 7B). As a reference, the median MAPK activity of cells imaged under the same conditions but in absence of a mating partner is plotted. In the fusing cell, a gradual increase in MAPK activity starts 40 to 60 minutes prior to fusion. In parallel, we observe that the median CDK activity is low for all time-points. However, the 75 percentile stabilizes to a low value 40 minutes prior to fusion, denoting that a fraction of the population enters in G1 within the hour preceding the fusion.

From the analysis of these fusion events, it becomes clear that the enrichment in G1 state prior to fusion occurs in cells that experience a low level of MAPK activity, or, in other words, when cells are surrounded by a low concentration of mating pheromone. In the exogenous stimulation experiments, we have shown that at low pheromone concentrations, a fraction of the cells do not commit to the mating response and keep proliferating. We verified if in the mating mixtures non-fusing cells were influenced by the presence of the mating partners. To achieve this, we monitored the CDK activity in cells that were cycling through G1. Interestingly, we observe that the G1 state of these cells is prolonged and displays a great variability (Figure 7C). In these cycling cells, the CDK activity remains lower in cells in mating mixtures compared to the same cells imaged alone (Figure 7D). In parallel, a weak activation of the mating pathway can be observed during the G1 phase (Figure 7E). *ste5*Δ cells imaged in presence of a mating partner display a fast cycling through G1 compared to the WT cells because they remain insensitive to the presence of the partners. In this background, the deletion of the scaffold Ste5 leads to an absence of activity of the MAPK Fus3 and Kss1 which cannot counteract the rise of the CDK activity at START.

## Discussion

In this study, we performed dynamic single cell measurements with live-cell imaging. These time lapse movies were automatically quantified, allowing the clustering and *in silico* synchronization of hundreds of single cell traces. The ability to follow the response of individual cells in a population of naturally cycling cells has enabled us to monitor the influence of the cell cycle on the mating process with minimal perturbations. Importantly, the correlation of multiple signaling activities within the same cell by combining fluorescent reporters for CDK and MAPK activities allowed us to identify different MAPK signaling patterns, which demonstrates the ability of the MAPK cascade to integrate the CDK activity to deliver the required signaling output. The molecular mechanisms of some of these integrations have been identified previously, but our quantitative measurements suggest that additional mechanisms contribute to the interplay between MAPK and CDK.

We have verified with exogenous stimulation experiments performed with various concentrations of pheromone that START is a central integration point where cells compare the relative levels of MAPK and CDK activities to decide on their cellular fate: proliferation or mating. It remains to be precisely determined which molecular mechanisms control this decision. On the MAPK side, our data agree with the numerous previous studies that demonstrated that the MAPK activity controls the CDK via the protein Far1 (Peter *et al*., 1993; Peter and Herskowitz, 1994). The mating pathway regulates both the level of Far1 and its phosphorylation status. What remains to be understood is how each one of these changes influence the decision made by the cell.

On the CDK side, Cdc28 kinase activity is directly regulating the signal flow in the MAPK pathway. Blocking Cdc28 activity relieves this inhibition allowing recovering full MAPK activity, within minutes after addition of the chemical inhibitor (Figure 5). This suggests a very direct mechanism of action on the mating pathway. The primary candidate for this process has been Ste5. However, our quantitative measurements speak for an additional mechanism detectable mostly at low pheromone concentrations. Other potential candidate targets of Cdc28 could be the G-protein and the receptor, or other proteins in the MAPK signal transduction cascades.

The experiments performed with mating mixtures also illustrate that the interplay between the CDK and the MAPK are central elements in the cell fate selected by the cells. Our data indicate that the commitment to the mating program is taking place at low levels of MAPK signaling activity, where we have shown with exogenous pheromone stimulation, that the CDK can frequently override the MAPK arrest. The sensing of the pheromone secreted by nearby mating partners triggers a low level of MAPK activity, including the activation of positive and negative feedback regulation mechanisms (Figure 8). In the cells that will successfully mate, we observe that the MAPK activity is progressively rising and thereby preventing an activation of the CDK that would promote an entry in the cell cycle. The polarized secretion of pheromones by both partners contributes further to this increase in MAPK activity that reaches its maximum shortly before the fusion of the two cells. Cells which are at some distance from a potential partner experience lower levels of pheromone. Internal feedback loops together with the secretion of the Bar1 protease contribute to dampen the MAPK activity thus when cells reach the cell-cycle commitment point the CDK can override this low mating signaling activity and promote a new cell cycle round.

**Figure 8:**
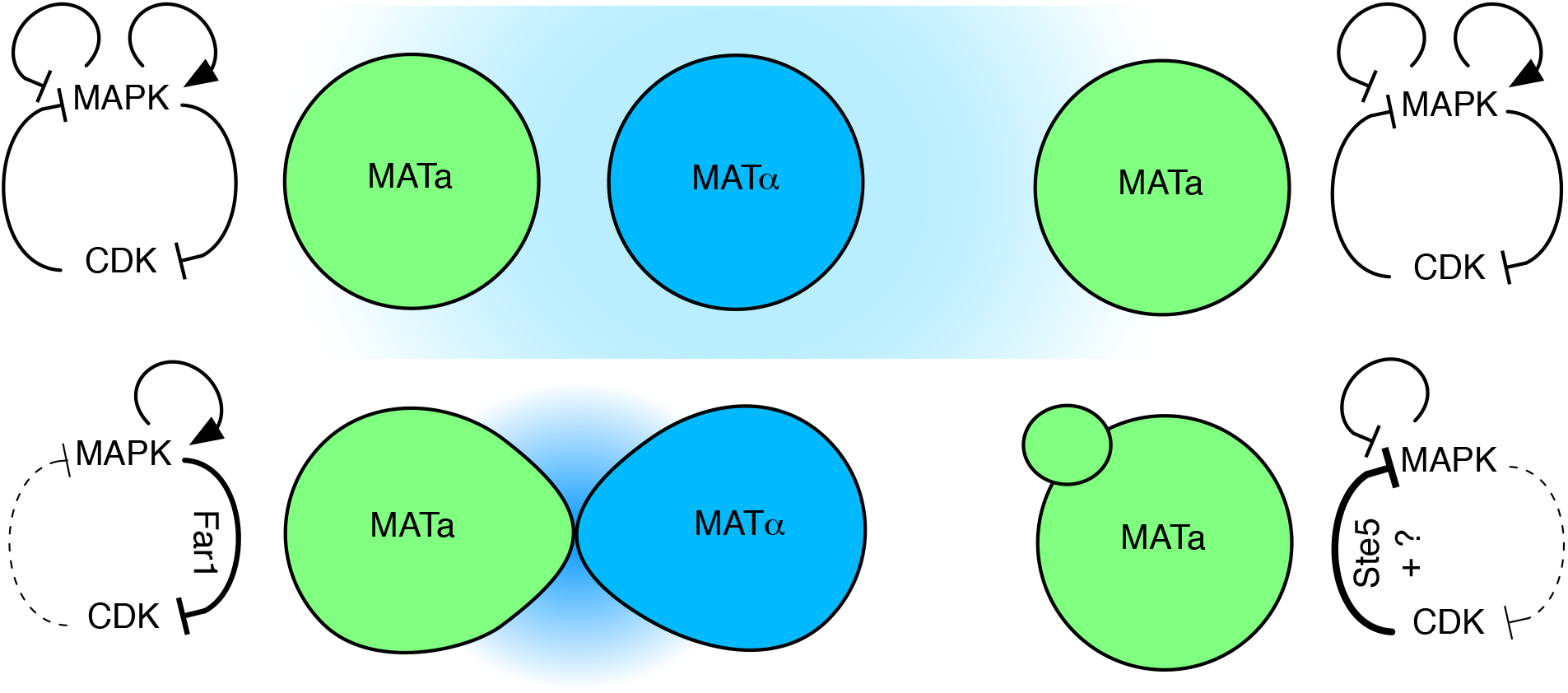
Scheme of the interplay between CDK and MAPK during mating. *MAT***a** cells respond to a pheromone gradient secreted by a *MAT***α** mating partner. The strength of the MAPK activity will depend on the level of α-factor sensed by the cells and the contribution of positive and negative feedback. At START, relative CDK and MAPK activities will be compared in order for the cell to select a fate: division or mating.

Throughout all eukaryotes, MAPK signaling and the cell cycle are highly conserved cellular processes. The homolog of Fus3 in mammalian cells is ERK. ERK is probably best known for its implication in cell growth and division in somatic cells (Meloche and Pouysségur, 2007) and hyperactivating mutation in the ERK pathway are found in numerous cancers (Davies *et al*., 2002). However, the ERK pathway can also inhibit cell cycle progression. One well-known example is the stimulation of PC-12 cells with NGF, which results in a prolonged activation of ERK, promoting a cell cycle arrest and differentiation into neuronal cells (Marshall, 1995; Pumiglia and Decker, 1997). More generally, during development MAPK pathways play a central role in the commitment of cells into specific lineage and this is often performed in tight correlation with the cell cycle (Orford and Scadden, 2008). Correlating dynamic measurements of CDK and MAPK activities in these cells could reveal the underlying mechanisms that allow the MAPK to integrate cell cycle cues to modulate the signaling outcome.

## Materials and Methods

### Strains and plasmids

Yeast strains and plasmids are listed in Supplementary Tables 1 and 2. SKARS plasmids were constructed by cloning the different sensors from Durandau et al (Durandau *et al*., 2015) into a pSIV vector backbone (Wosika *et al*., 2016). The sensor and the corrector were assembled by cloning the *STE7_1-33_-NLS-NLS-mCherry* or the *STE7_ND_-NLS-NLS-CFP* (HindIII-KpnI) sequences downstream the p*RPS6B* promoter into the pSIV-URA (SacI-KpnI). The plasmid pED141 containing both sensor and corrector was obtained by cloning the p*RPS6B -STE7_ND_-NLS-NLS-CFP* into the second MCS of pED92 (AatII-SphI). Plasmids were transformed in yeast of W303 background expressing the Hta2-iRFP (yED136) or the Hta2-tdiRFP (yED152). Whi5 was tagged with mCitrine using pGTT-mCitrine plasmid (Wosika *et al*., 2016). Kss1 was tagged with mScarlet (pGTL-mScarlet). p*AGA1* induction was monitored by integrating the p*AGA1*-dPSTR^R^ in the URA3 locus (Aymoz *et al*., 2018).

Genomic deletions were constructed in cells bearing the sensors using KAN or NAT resistance cassettes (Longtine *et al*., 1998; Goldstein and McCusker, 1999). Plasmids containing the coding sequence of Ste5, Ste20 and Ste7 were obtained by amplification of a chromosome fragment spanning from −100bp (into the promoter) to the end of the ORF and cloned using PacI-NheI sites into the pGTH-CFP (replacing the fluorescent protein). Mutated variants were then obtained by replacing a portion of the plasmid DNA coding sequence by a synthetic double-stranded DNA fragment bearing the desired mutations. Plasmids were integrated into the genome of yeast by replacing the NAT or KAN cassettes used to delete the native gene using homology regions in the promoter of the gene and in the TEF terminator of the deletion cassette, which is also present on the pGTH vector.

### Sample preparation

The cells were grown to saturation in overnight cultures at 30°C in synthetic medium (YNB: CYN3801, CSM: DCS0031, ForMedium). They were diluted in the morning to OD_600_=0.05 and grown for at least four hours before starting the experiments. For experiments in wells, 96-well plates (MGB096-1-2LG, Matrical Bioscience) were coated with a filtered solution of Concanavalin A (0.5 mg/ml, 17-0450-01, GE Healthcare) for 30min, rinsed with H_2_O, and dried for at least 2 hours. Before the experiment, the cultures were diluted to an OD_600_ of 0.05, and briefly sonicated. A volume of 200μl of culture was loaded into each well. Cells were left settling 30–45 minutes before imaging. To stimulate the cells, 100μl of inducing solution was added to the wells. For α-factor stimulation, final concentration is indicated into the figure legend. To inhibit the *cdc28-as*, a 25mM stock solution of NAPP1 (A603004, Toronto Research Chemical) in DMSO was diluted in SD-medium and added to the wells. The final working concentration is indicated in the figure legend. For control experiments, cells were treated with DMSO 0.16% in SD-full.

For mating assay experiments, log phase cultures of *MAT***a** and *MAT***α** (ySP694) were diluted to OD 0.1 in 500μl SD-full. Cells were spun down for 2min at 3000rpm. The supernatant was removed from the cultures and *MAT***α** cells were resuspended in 10μl of SD-full. These 10μl were then used to resuspend the *MAT***a** cells. 1μl of this mating mixture was placed on an agar pad. The pad was then placed upside down into the well of a 96-well plate. To prepare the pad, a 2% agarose mixture in SD-full was heated for 5 min at 95°. 150μl liquid was placed in a home-made aluminum frame. After cooling down, the 7 mm square pad was gently extracted from the frame. Typically, 6 to 8 mating pads were prepared and imaged in parallel.

### Microscopy

Images were acquired with a fully automated inverted epi-fluorescence microscope (Ti-Eclipse, Nikon), controlled by Micromanager software (Edelstein *et al*., 2010) and placed in an incubation chamber set at 30 °C, with a 40X oil objective and appropriate excitation and emission filters. The excitation was provided by a solid-state light source (SpectraX, Lumencor). The images were recorded with a sCMOS camera (Flash4.0, Hamamatsu). A motorized XY-stage allowed recording multiple fields at every time points. Cy5p5 (200ms), CFP (100ms), RFP (100ms), YFP (300ms) and two brightfield (10ms) images were recorded at time intervals of 3 or 5 minutes.

### Data analysis

Time-lapse movies were analyzed with the YeastQuant platform (Pelet *et al*., 2012). The cell nuclei were segmented by thresholding from Cy5p5 images. The contour of the cell around each identified nucleus was detected using both brightfield images. The cytoplasm object was obtained by removing the nucleus object expanded by two pixels from the cell object. The nuclei were tracked across all the frames of the movie. Multiple features of each object were quantified. Note that for mating pad experiments, the *MAT***α** cells were not segmented as they do not express the Hta2-iRFP. Dedicated scripts were written in Matlab R2016b (The Mathworks) to further analyze the data. Except for Figure 7, only cells tracked from the beginning to the end of the movie were taken into consideration. In addition, a quality control was applied on each trace and only the traces with low variability in nuclear and cell area, and Cy5p5 nuclear fluorescence were kept for further analysis. On average, 65% of the cells tracked from the beginning to the end of the time lapse passed the quality control.

For each cell, the average nuclear intensity in the fluorescent channel corresponding to the SKARS, the Corrector and Whi5-mCitrine were divided by the average intensity in the cytoplasm for every time point (ratio N/C). To quantify the MAPK activity, the Adjusted ratio for each individual cell was obtained by dividing the ratio N/C of the Corrector by the ratio N/C of the SKARS. The CDK activity of each cell was defined as the inverse of the Whi5-mCitrine N/C ratio. The initial CDK activity was defined as the average of three time-points before the stimulus (3–6 minutes before α-factor stimulus). All these quantities are unitless numbers, since they are ratios between fluorescence intensities obtained from the microscope camera. We used the symbol [–] to represent this lack of units.

### CDK activity-based clustering and synchronization

When plotting a histogram using the Whi5 ratio N/C (Nuclear intensity over Cytoplamsic intensity) of all cells at all time points, we identified two populations. To properly distinguish between ratios corresponding to nuclear Whi5 versus ratios corresponding to cytoplasmic Whi5, we first separated ratio values using a fixed threshold (threshold=2). However, this value overestimated the limit of what could be considered as nuclear Whi5. We then identified cells with ratios below this threshold at all time points and replotted a histogram of all Whi5 ratio N/C values from this sub-population. This method enables to enrich the sample with low N/C ratio. This second histogram presented a clearer overview of the values corresponding to cytoplasmic Whi5. We then calculated the derivative of the histogram and identified the position of fifth-lowest derivative value. The corresponding Whi5 ratio N/C was used as a final threshold to separate nuclear to cytoplasmic Whi5 values. This procedure enables to correct for slight differences between experiments mostly due to the chosen experimental settings (96 well-plate or pad experiments). Globally, the procedure adjusted the threshold between 2 and 1.7. If the Whi5 ratio N/C was above the threshold, corresponding to a low CDK activity value (C/N) and thus the cell was considered in G1. On the opposite, values below the threshold indicate that the cell was *Out-of-G1. G1* cells sub-population was defined by single cell traces which remained above the threshold for the entire duration of the time lapse. Conversely, *Out-of-G1* cells cluster was defined by traces that stay below this threshold. The other single cell traces are scanned for G1-entry and G1-exit events. If a pattern of two values below the threshold followed by two values above the threshold was found, we considered that the cell belonged to the *G1-entry* cluster. The same strategy was used to identify the *G1-exit* sub-population, using a pattern of two values above the threshold followed by two values below the threshold. Cells in the *G1-entry* cluster were defined as having a single event of G1-entry. Cells in the *G1-exit* cluster were selected as having only one event of G1-exit. Finally, the *Through-G1* cluster is made of cells that display a G1-entry event followed by a G1-exit event. Because the duration of the time-lapse experiments is much shorter than the cell cycle of the yeast (45 min vs. 90 min), we considered that the number of events cannot exceed two. Cells which do not follow this criterion were rejected from the analysis. Depending on the experiment, we rejected a maximum of 5% of the total population. The position of the G1-entry was used to create new time vectors for synchronization of the *G1-entry* and the *Through-G1* single cell traces. We extracted the individual time vector for each cell and subtracted the time at which the G1-entry event occurred.

### Detection of fusion events in mating assays

*MAT***α** cells express the CFP fluorescent protein under the control of a strong promoter. When *MAT***a** and *MAT***α** cells fuse the CFP from the *MAT***α** cell diffuses into the partner leading to a rise of the fluorescence into the segmented *MAT***a**. Fusing *MAT***a** cells were first detected bioinformatically by identifying a sudden increase of the CFP intensity in the nucleus object. We chose the nucleus object rather than the whole cell to avoid that mis-segmentation triggers a misleading increase in CFP intensity not linked to a fusion event. Fusion event time was defined as the time point at which the difference in the CFP nuclear signal between two consecutive time points exceed 200, while remaining below 700. Traces of fusing cells were selected as they contain a single detected fusion event. We filtered out cells with aberrant absolute and derivative nuclear CFP fluorescence signal or tracked for less than 15 min prior or after the fusion time. CFP images corresponding to selected cells were checked manually to reject false positives that could not be detected during the bioinformatics analysis. The position of the fusion event into the time vectors of individual fusing cells was used as a time reference to align temporally the MAPK activity and CDK (Figure 7).

## Acknowledgments

We thank all members of the Pelet and Martin labs for helpful discussions and comments on the manuscripts, Marta Schmitt and Clémence Varidel for technical assistance. This study was supported by Swiss National Science Foundation grants (PP00P3_139121) and the University of Lausanne.

## Author contributions

ED and SP conceived and study and wrote the manuscript. ED constructed the strains and performed the experiments with exogenous stimulation of pheromone. SP performed the mating assay experiments. ED analyzed the single cell data.

The authors have declared that no competing interests exist.

## Supplementary Material

Supporting Information Legends

Supplementary Figures 1 to 11

Supplementary Tables 1 and 2

