## Supplementary Tables 1 and 2 and Supplementary Figures 1 to 11 for "Cross-regulation between CDK and MAPK control cellular fate"

**Eric Durandau, Serge Pelet**
Department of Fundamental Microbiology, University of Lausanne, CH-1015 Lausanne, Switzerland

**Supplementary Figures 1 to 11**

**Supplementary Tables 1 and 2**

### Supplementary Figure Legends

**Supplementary Figure 1. Improvement of the MAPK activity quantification.**
**A, D.** Synchronized CDK activity signals. CDK activity of cells entering in G1 plotted relative to time of strongest decrease in CDK signal (see also Supplementary Fig 3). **B, E**. Dynamic subcellular localization of the SKARS and Corrector in cells entering in G1. SKARS and Corrector nuclear over cytoplasmic ratio (N/C) of cells entering into G1 were measured. Adjusted ratio was obtained by dividing the Corrector N/C by the SKARS N/C. For comparison, SKARS N/C, Corrector N/C and Adjusted ratio were normalized relative to their respective mean and then plotted relative to the time of G1 entry as in A and D. **C, F**. Normalized standard deviation of SKARS N/C, Corrector N/C and Adjusted ratio along the time axis calculated for all cells entering G1 and plotted as boxplot. All the analysis was performed from the time lapse experiment presented in Figure 1C.

**Supplementary Figure 2: Synchronization of CDK activity traces relative to G1.**
WT cells expressing SKARS and CDK sensors were imaged for 2 h with five-minute intervals. Images were analyzed and the CDK activity was quantified as the ratio of yellow fluorescent intensity between the cytoplasm and the nucleus for each individual cell and at each time point. Single cell traces that displayed a drop and rise in CDK activity were selected (Nc = 70). **A.** Single cell traces and the median of the population are plotted relative to the real experimental time where division happens in an asynchronous manner. **B.** The same single cell traces are synchronized relative to G1 (CDK activity minimum set as time 0). The median of these synchronized traces displays a sharp trough around time zero and lower ones at -90min and +90min matching well the cell cycle length.

**Supplementary Figure 3. CDK and MAPK activities relative to G1-entry.**
WT cells expressing SKARS and Whi5 sensors were imaged for 45 minutes with three-minute intervals. 1μM α-factor was added to the well nine minutes after the acquisition start **A.** The median CDK (green) and MAPK (red) activities of cells from the *G1-entry* cluster are plotted relative to the time of α-factor addition (solid lines) along with a few single cell traces (dashed lines). **B.** The last time point at which the CDK activity remains above 0.55 is taken as a temporal reference and used to align temporally the single cell traces. CDK (green) and

MAPK (red) median activities and the same single cells as in panel A are plotted relative to this reference time.

**Supplementary Figure 4. Dynamic MAPK activity of cells transiting through G1.** Experiments from Figure 2 are analyzed using the clustering strategy described in the Methods. Cell transiently passing through G1 during the time lapse experiments are identified as they exhibit a trough in CDK activity. Single cell traces were synchronized with the same strategy used for the G1 entry population. **A.** CDK (green) and MAPK (red) activities from the experiment presented in Figure 2A-D. **B.** The dynamic MAPK activity of at least three replicates for different pheromone concentrations is averaged and plotted as in Figure 3.

**Supplementary Figure 5. Monitoring mating pathway activity with Kss1 relocation and p*AGA1*-dPSTR$^{R}$ reporters**
**A.** Scheme of the Kss1 relocation mechanism. Under vegetative growth conditions, Kss1 is anchored in the nuclei of the cells via its association with Ste12, Dig1 and Dig2. Disruption of this complex by MAPK activity results in a homogenous distribution of Kss1 throughout the cell. **B.** Cells bearing the Kss1-mScarlett and Whi5-mCitrine cell-cycle reporter are stimulated at time 0 with pheromone. **C.** Scheme of the dPSTR assay. Activation of the p*AGA1* promoter upon mating pathway stimulation induces the expression of a small peptide (SZ-NLS: SynZip – Nuclear Localization Sequence). This small peptide has a strong affinity for an endogenously expressed fluorescent protein functionalized with a compatible SynZip. The strong heterodimer formed between the SynZip results in an accumulation of the fluorescence in the nucleus of the cell. **D.** Cell bearing the p*AGA1*-dPSTR$^{R}$ system and Whi5 tagged with mCitrine are imaged upon stimulation with pheromone at time 0.

**Supplementary Figure 6. Dynamics of MAPK activity monitored by Kss1 relocation in different cell cycle phases**
**A-D.** Kss1 relocation (cytoplasmic over nuclear fluorescence) was quantified for cells in an asynchronously growing culture stimulated with 1μM α-factor at time 0. 228 single cell traces were clustered by comparing the CDK activity prior and after stimulation (see Methods and Figure 2). *G1* cells (A, Nc =83) *Out-of-G1* cells (B, Nc =92) *G1-exit* cells (C, Nc =16) and *G1-entry* cells (D, Nc =35) were clustered based on their CDK activity pattern. The median (solid line) Kss1 relocation (red) and CDK activity (green) and the 25-75 percentiles (shaded area) of each sub-population are plotted.
**E.** Summary of the various dynamic Kss1 relocation behaviors observed in the four main cell cycle clusters. Each line corresponds to the mean of the medians of 4 biological replicates. Error bars represent the standard deviations of the medians.

**Supplementary Figure 7. Dynamics of mating pathway transcriptional activity monitored by the p*AGA1*-dPSTR$^{R}$ relocation in different cell cycle phases**
**A-D.** p*AGA1* promoter activity monitored by the dPSTR relocation (nuclear minus cytoplasmic fluorescence) was quantified for cells in an asynchronously growing culture stimulated with 1μM α-factor at time 0. 419 single cell traces were clustered by comparing the CDK activity prior and after stimulation (see Methods and Figure 2). *G1* cells (A, Nc =80) *Out-of-G1* cells (B, Nc =40) *G1-exit* cells (C, Nc =56) and *G1-entry* cells (D, Nc =213) were clustered based on their CDK activity pattern. The median (solid line) dPSTR relocation (red) and CDK activity (green) and the 25-75 percentiles (shaded area) of each sub-population are plotted.
**E.** Summary of the various dynamics of p*AGA1* induction observed in the four main cell cycle clusters. Each line corresponds to the mean of the medians of 4 biological replicates. Error bars represent the standard deviations of the medians.

**Supplementary Figure 8. Analysis of MAPK activity and corrector nuclear enrichment in different cell cycle clusters.**
**A.** Median and 25-75 percentile of the *G1*, *Out-of-G1*, *G1-exit* and *G1-entry* clusters are plotted for the MAPK activity, CDK activity and N/C (Nuclear to Cytoplasmic ratio) of the Corrector signal. For *G1*, *Out-of-G1*, *G1-exit*, time 0 corresponds to the addition of pheromone. For the *G1-entry* cluster, traces were aligned respective to the time of entry into G1 based on the CDK signal. **B.** Initial N/C ratio of the corrector for the whole population or the *G1* and *Out-of-G1* cluster. The vertical dashed line represents the threshold used to filter the cells for low or high corrector enrichment. **C.** Correlation between the initial nuclear to cytoplasmic ratio in the sensor and the corrector channels. **D.** and **E.** Similar graphs as in panel A for cells whose corrector N/C remains below (D) or above (E) the threshold throughout the time lapse.

**Supplementary Figure 9. Pheromone dose-dependent MAPK activation in BAR1 background.**
Experiments performed in Figure 3 were reproduced with the BAR1+ strain. **A.** Population level MAPK activity in BAR1 cells following stimulation with various α-factor concentrations **B-D**. Dynamic MAPK activity for three clusters described in Figure 2. In panels A to D, each line is the mean of at least three replicates medians at a given α-factor concentration. Error bars represent the standard deviations. **E.** Fractions of cells in the various cell cycle sub-populations as function of pheromone concentration in the BAR1 strain. Values are means of at least three replicates.

**Supplementary Figure 10: MAPK-dependent cell cycle arrest in G1 requires Far1.**
**A.** Pheromone dose-dependent activation of the MAPK in WT (red) and *far1Δ* (blue) clustered based on CDK activity pattern. The MAPK response is defined as the median MAPK activities at three to five minutes post-stimulation minus the median MAPK activity pre-stimulation. Error bars are standard deviations of at least three replicates.
**B.** Fractions of cells accumulated in G1 at the end of time-lapse movies (sum of *G1* and *G1-entry* clusters) were plotted as function of the pheromone concentration. Values are the means of at least three replicates. Error bars represent the standard deviations.

**Supplementary Figure 11. Dynamic MAPK activity of cells expressing Ste20 or Ste7 mutants.**
**A, B.** Schematic representation of the various *STE20* (A) and *STE7* (B) alleles tested. The $Ste20_{11A}$ and $Ste7_{6A}$ variants are obtained by replacing the serine/tyrosine phosphorylation sites by an alanine (Ste20 positions: 269, 418, 423, 492, 502, 512, 517, 547, 555, 562, 573. Ste7 position: 105, 130, 167, 116, 137, 149). ND variants are obtained by replacing the coding sequences of the SLDDPIQF and the PLPPIPP motifs at protein positions 87 and 535 of Ste20 (A) or the SISLPPL motif at protein position 156 of Ste7 (B) with sequences of same length coding for alanines. Alleles are inserted at the original chromosomic locus and expressed under their native promoter into a strain bearing the biosensors and the $Ste5_{ND}$ allele. **C, D.** Dynamic MAPK activity of cells *Out-of-G1* cells and in *G1*. Cells are imaged and data are analyzed as in Figure 2. Lines correspond to the means of the median of at least three replicates. Error bars represent the standard deviations.

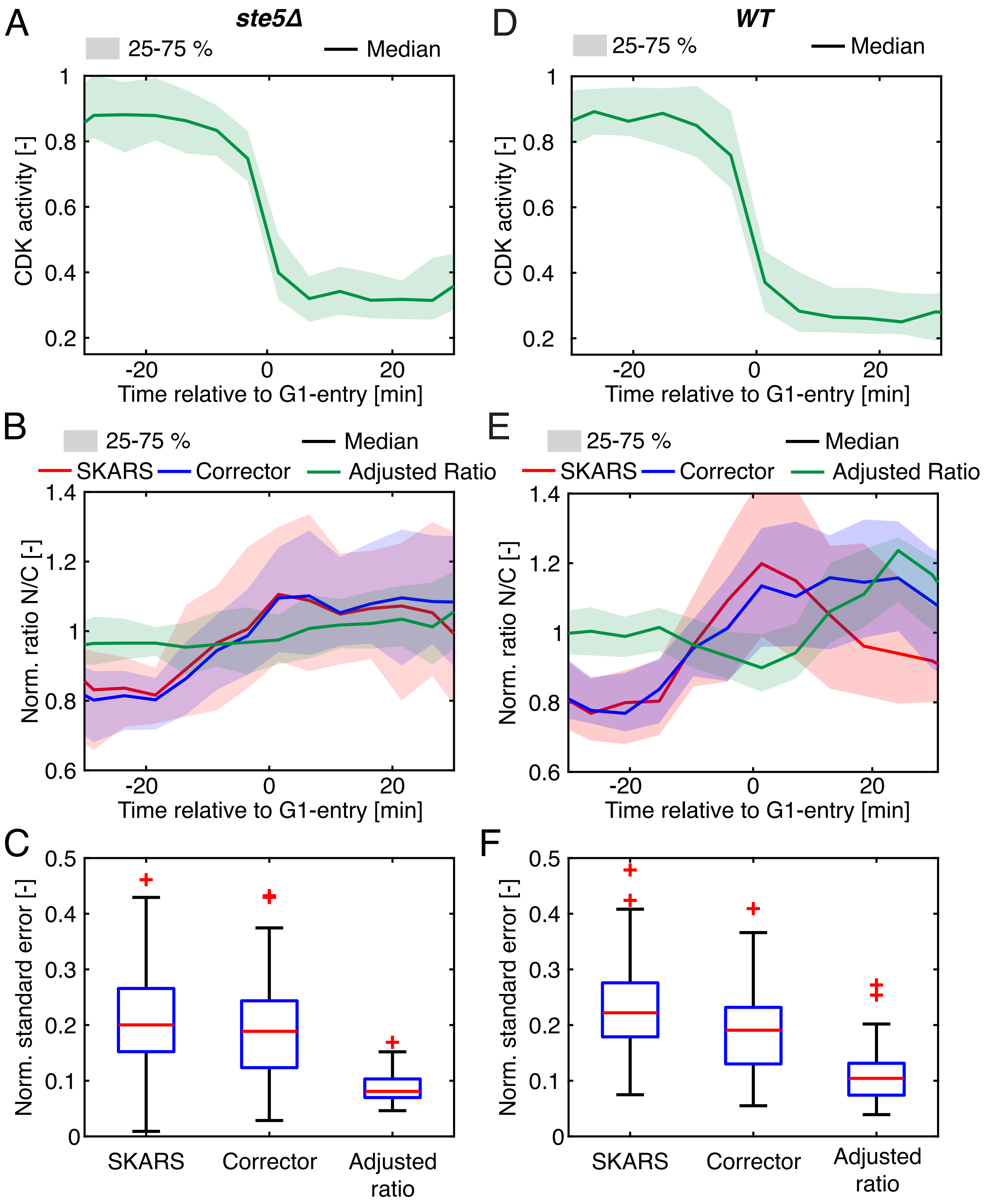


Supplementary Figure 1. Improvement of the MAPK activity quantification.

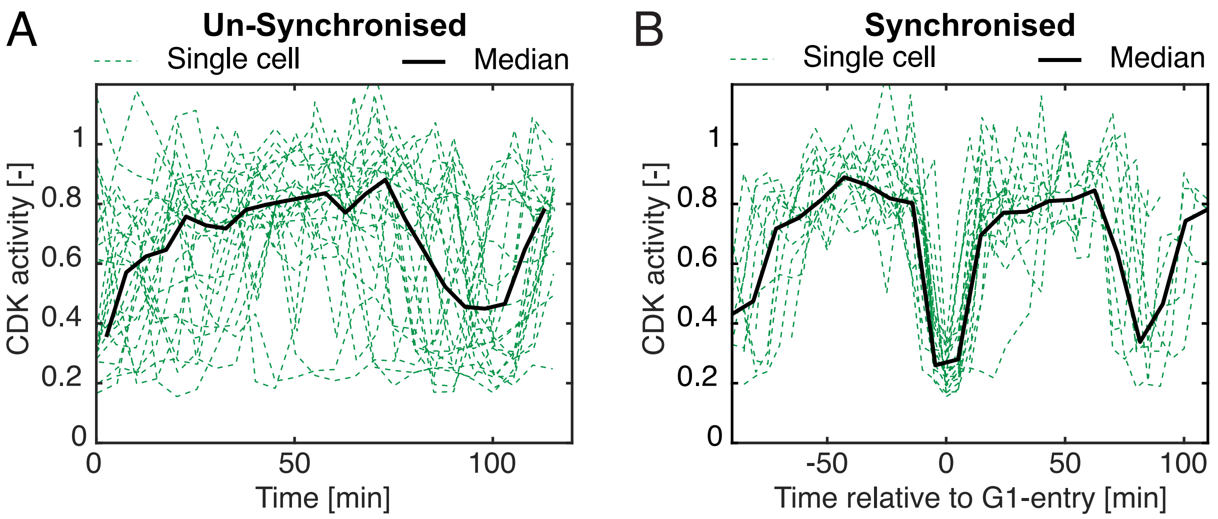


Supplementary Figure 2: Synchronization of CDK activity traces relative to entry into G1.

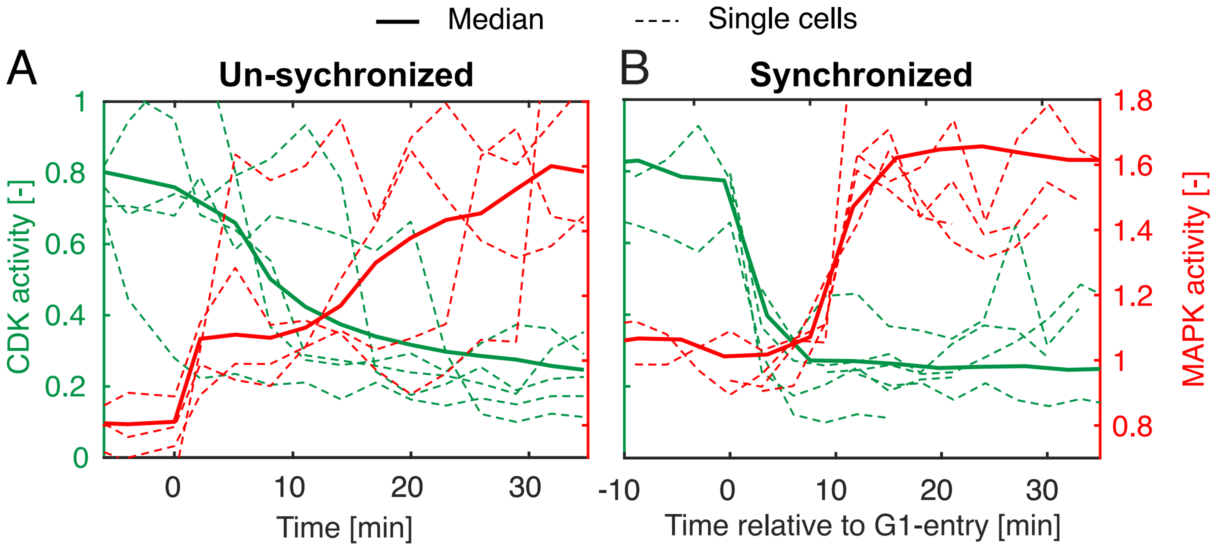


Supplementary Figure 3. CDK and MAPK activities relative to G1-entry.

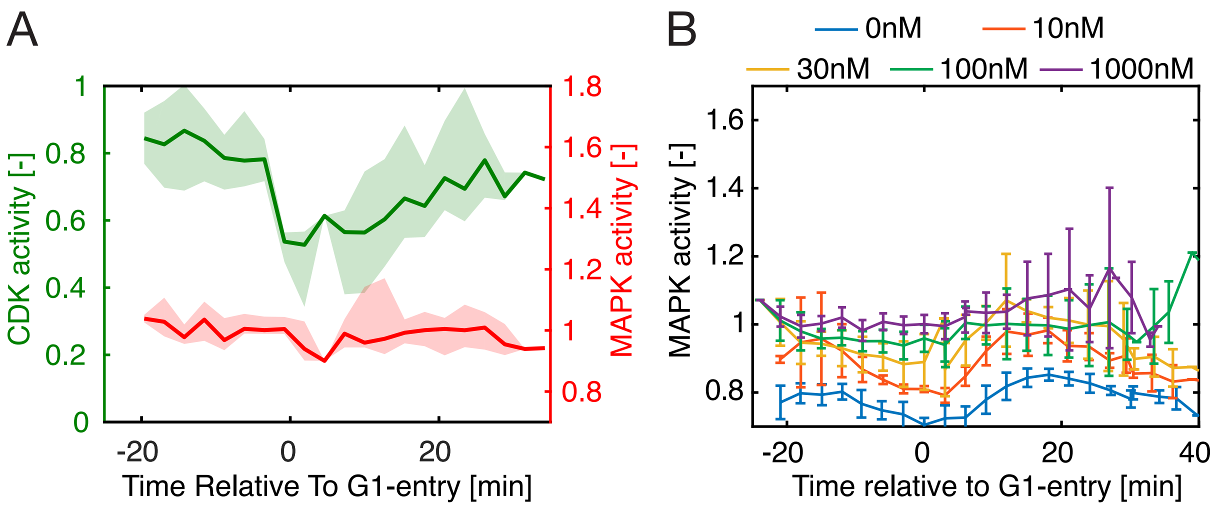


Supplementary Figure 4. Dynamic MAPK activity of cells transiting through G1.

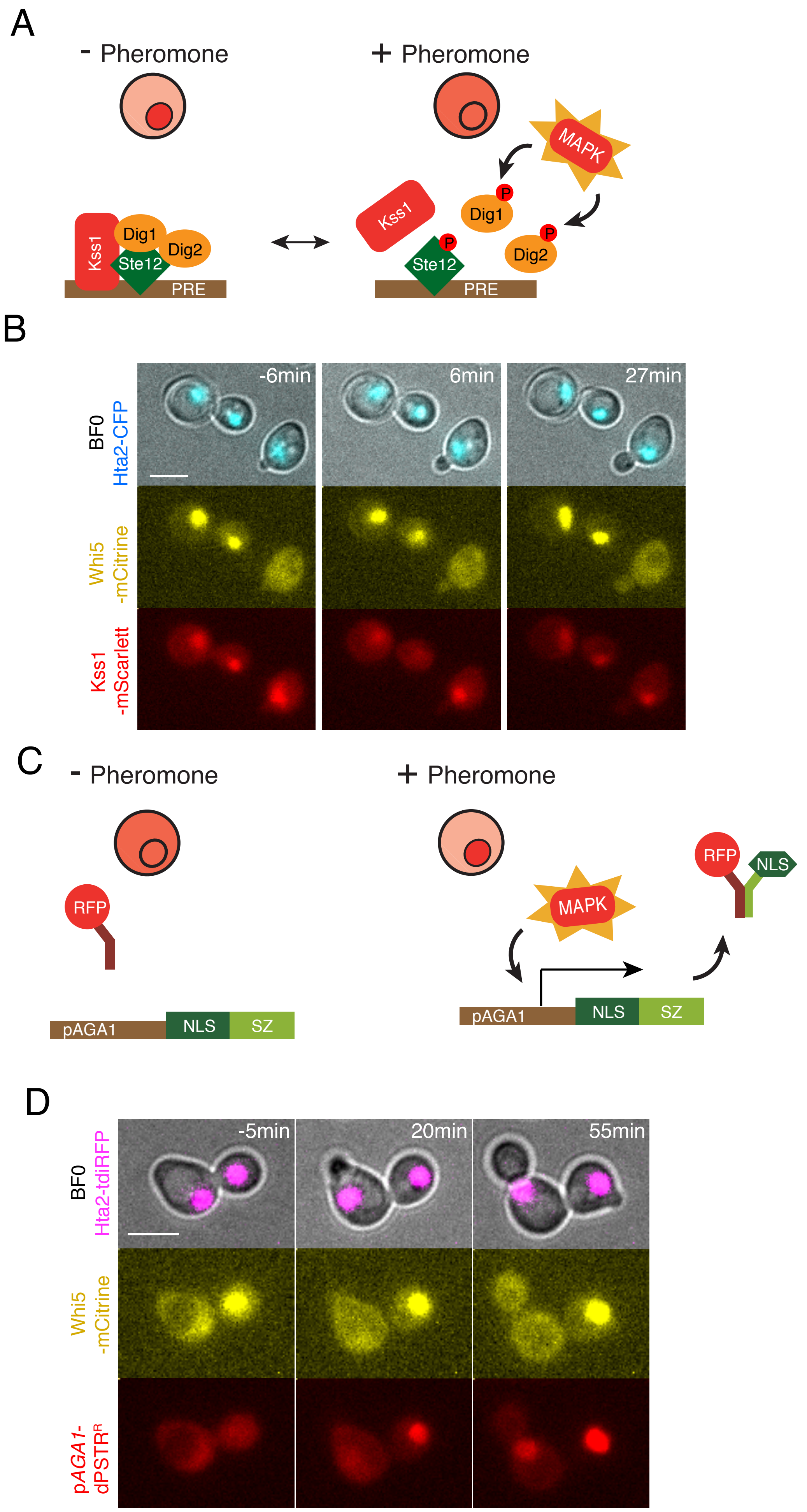


Supplementary Figure 5. Monitoring mating pathway activity with Kss1 relocation and p*AGA1*-dPSTR$^{R}$ reporters

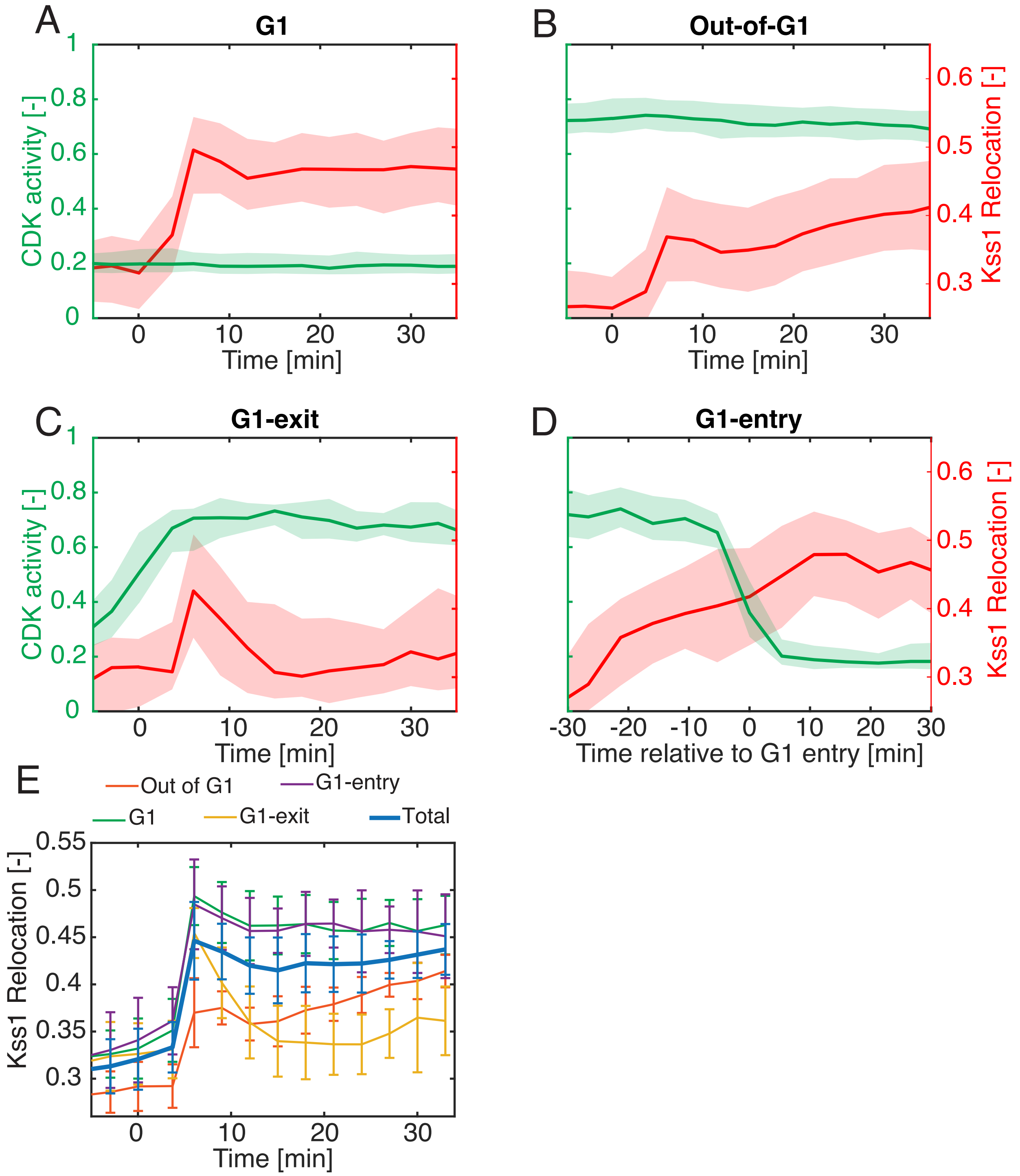


Supplementary Figure 6. Dynamics of MAPK activity monitored by Kss1 relocation in different cell cycle phases

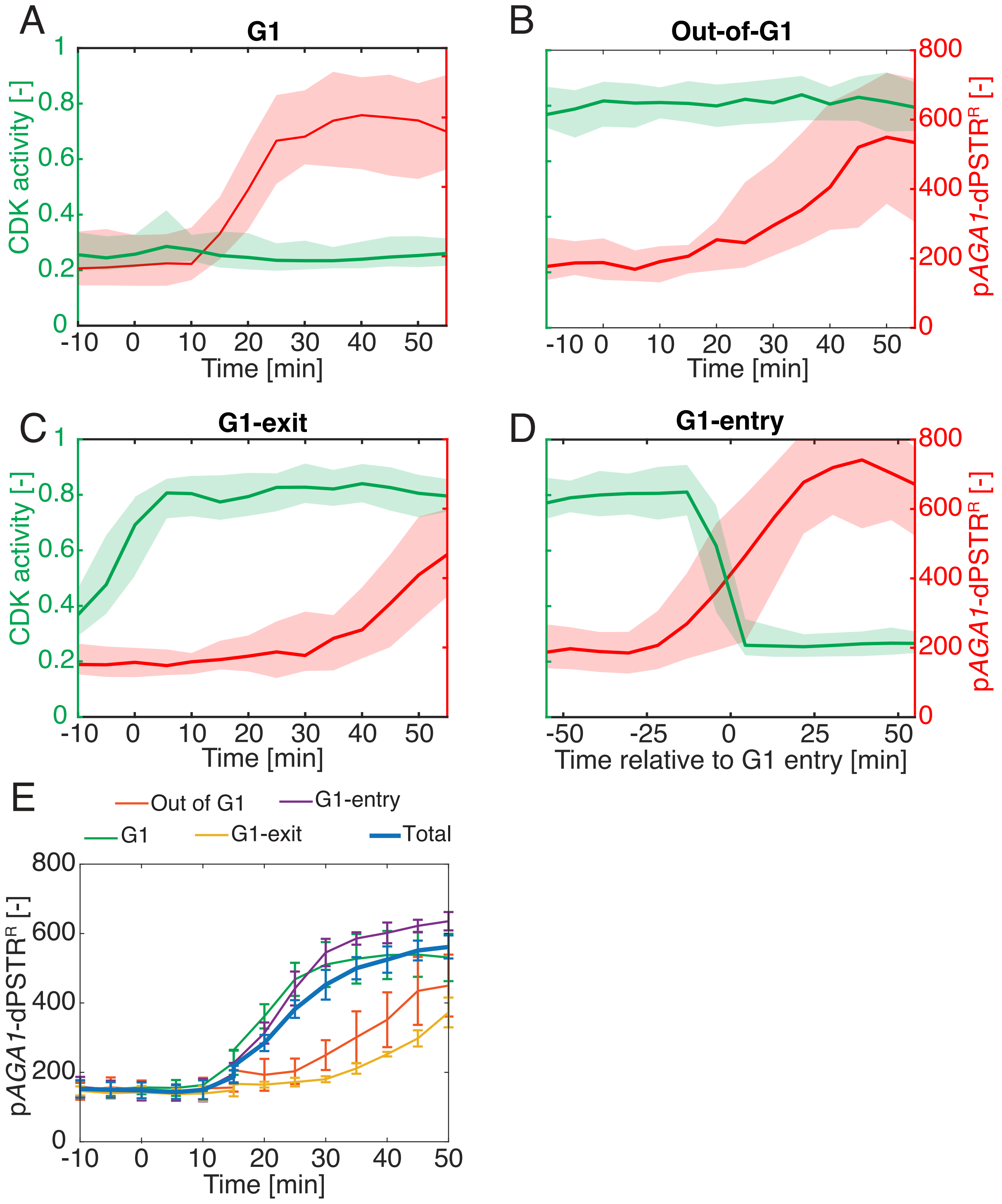


Supplementary Figure 7. Dynamics of mating pathway transcriptional activity monitored by the p*AGA1*-dPSTR$^{R}$ relocation in different cell cycle phases

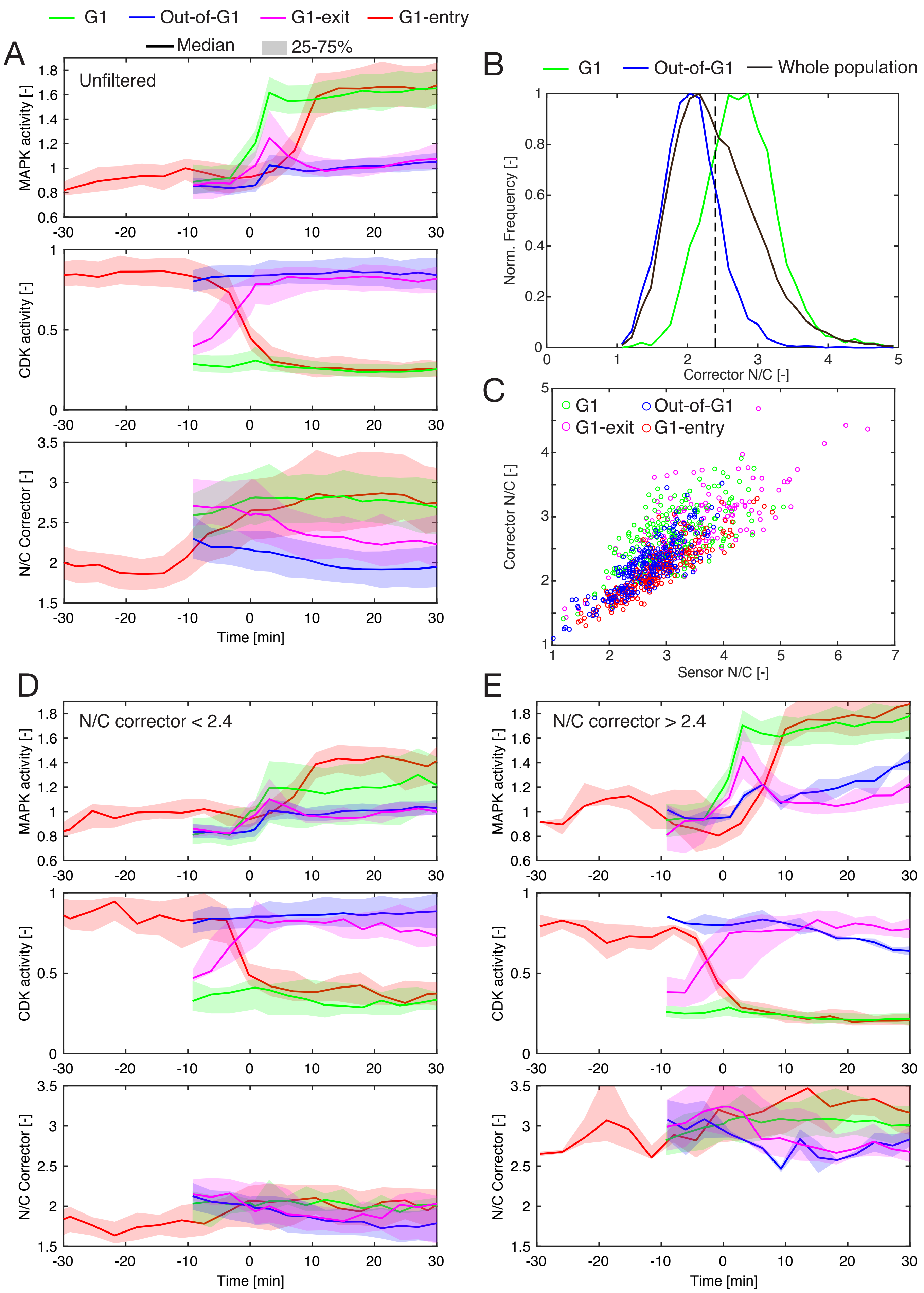


Supplementary Figure 8. Analysis of MAPK activity and corrector nuclear enrichment in different cell cycle clusters.

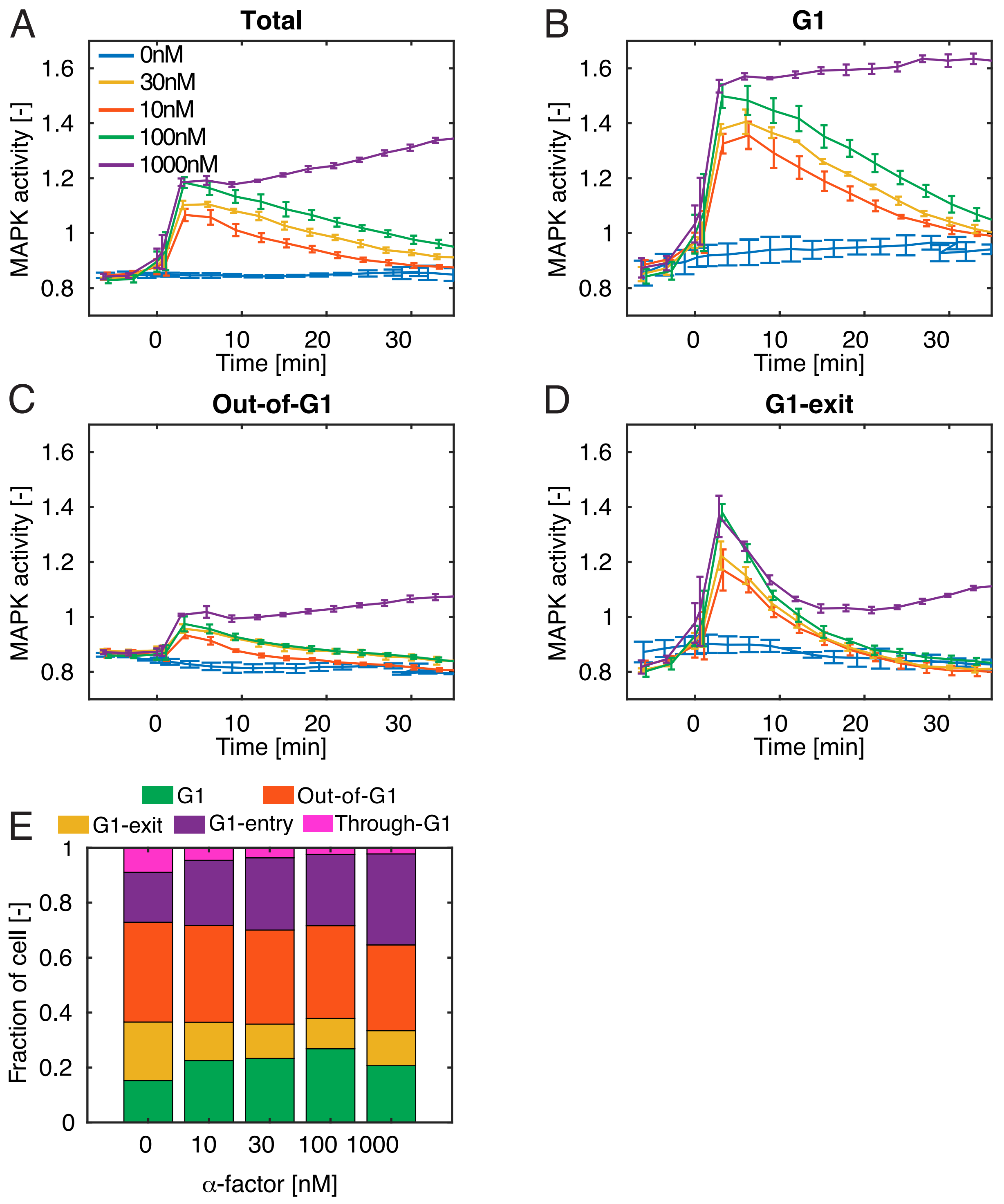


Supplementary Figure 9. Pheromone dose-dependent MAPK activation in BAR1 background.

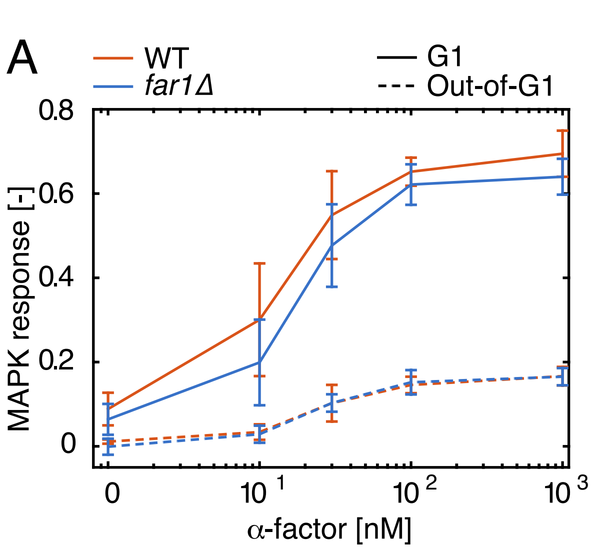


Supplementary Figure 10: MAPK dependent cell cycle arrest in G1 requires Far1.

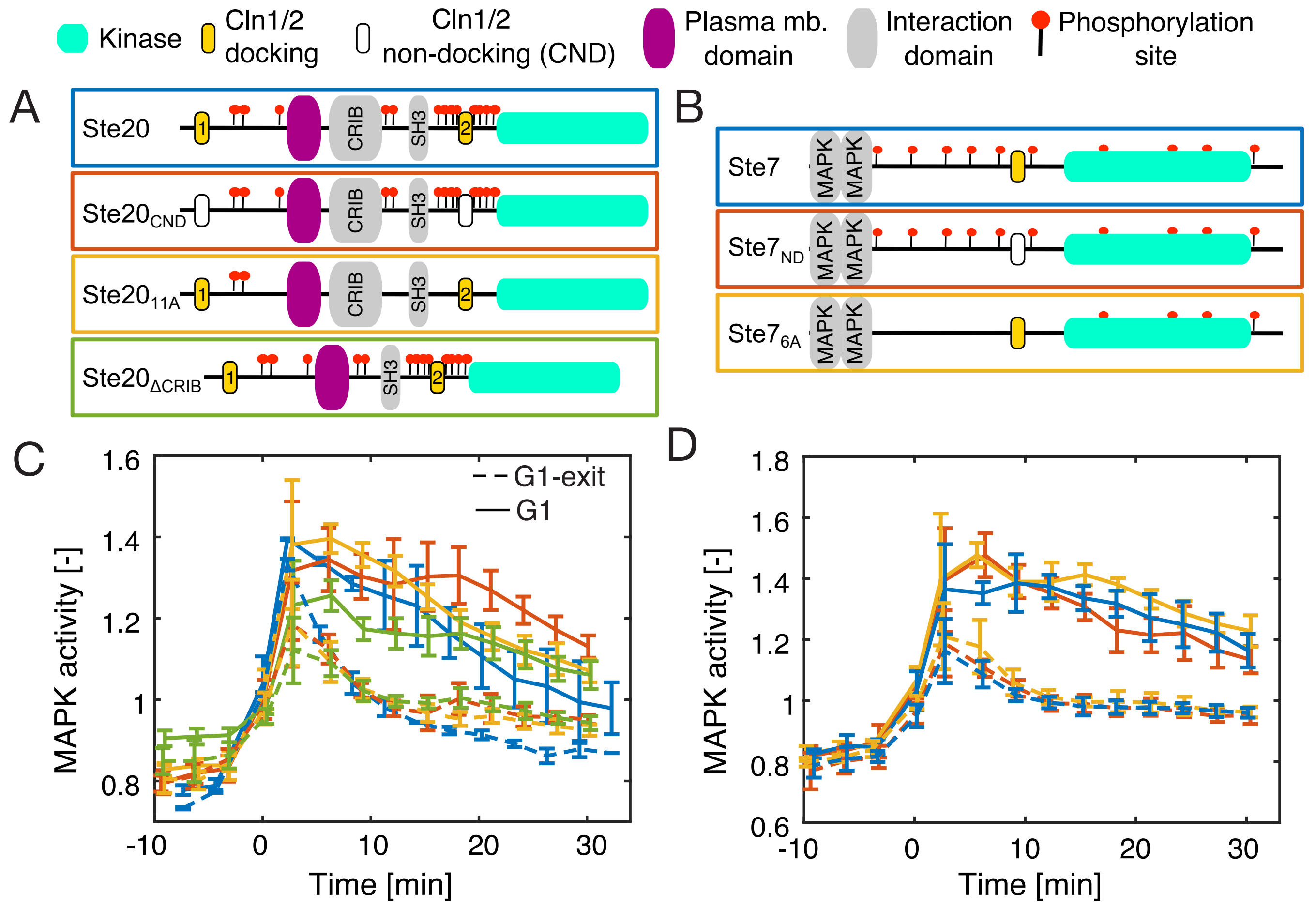


Supplementary Figure 11. Dynamic MAPK activity of cells expressing mutant allels of Ste20 or Ste7.

**Supplementary Table1: list of strains**

| Name | Genotype | Figure |
|---|---|---|
| ySP2 | *W303: MAT***a** *leu2-3, 112 trp1-1 can1-100 ura3-1 ade2-1 HIS3-11,15* | |
| ySP122 | *W303: MAT*$\alpha$ *leu2-3, 112 trp1-1 can1-100 ura3-1 ade2-1 HIS3-11,15* | |
| yED136 | *HTA2-iRFP-HIS* | |
| yED152 | *HTA2-iRFP-HIS ura3*:pED92 | |
| yED164 | *HTA2-iRFP-HIS ura3:*pED92 *leu2*:pED94 | |
| yED195 | $cdc28_{as}$ *HTA2-tdiRFP-HIS* | |
| yED210 | *HTA2-tdiRFP-HIS* | |
| yED238 | *HTA2-iRFP-HIS ura3*:pED92 *leu2*:pED94 *WHI5-mCitrine:TRP* | F1; S1-3 |
| yED253 | *HTA2-iRFP-KAN ura3*:pED92 *leu2*:pED94 *WHI5-mCitrine:TRP* ste5Δ::pED137 | F7 |
| yED255 | *HTA2-iRFP-HIS ura3*:pED92 *leu2*:pED94 *WHI5-mCitrine:TRP* *bar1Δ*::*NAT* | F2-4; S4, 10 |
| yED258 | *HTA2-iRFP-KAN ura3*:pED92 *leu2*:pED94 *WHI5-mCitrine:TRP* | |
| yED264 | *HTA2-iRFP-KAN ura3*:pED92 *leu2*:pED94 *WHI5-mCitrine:TRP* ste5Δ::*NAT* | F1, 7; S1 |
| yED270 | *HTA2-tdiRFP-HIS WHI5-mCitrine:TRP* | |
| yED271 | *HTA2-tdiRFP-HIS WHI5-mCitrine:TRP ste5Δ::NAT* | |
| yED275 | $cdc28_{as}$ *HTA2-tdiRFP-HIS WHI5-mCitrine:TRP* | |
| yED281 | $cdc28_{as}$ *tdHTA2-tdiRFP-HIS ura3*:pED141 *WHI5-mCitrine:TRP* | |
| yED284 | *HTA2-iRFP-KAN ura3*:pED92 *leu2*:pED94 *WHI5-mCitrine:TRP* ste5Δ::pED137 *bar1Δ*::*NAT* | F6, S8 |
| yED285 | *HTA2-iRFP-KAN ura3*:pED92 *leu2*:pED94 *WHI5-mCitrine:TRP* ste5Δ::pED138 *bar1Δ*::*NAT* | F6 |
| yED288 | $cdc28_{as}$ *HTA2-tdiRFP-HIS ura3*:pED141 *WHI5-mCitrine:TRP* *bar1Δ*::*NAT* | F5 |
| yED289 | *HTA2-iRFP-KAN ura3*:pED92 *leu2*:pED94 *WHI5-mCitrine:TRP* ste5Δ::pED147 | S11 |
| yED299 | *HTA2-iRFP-HIS ura3*:pED92 *leu2*:pED94 *WHI5-mCitrine:TRP* *bar1Δ*::*NAT far1Δ*::*KAN* | S11 |
| yED300 | *HTA2-iRFP-KAN ura3*:pED92 *leu2*:pED94 *WHI5-mCitrine:TRP* ste5Δ::pED147 *bar1Δ*::*NAT* | F6 |
| yED306 | *HTA2-iRFP-KAN ura3*:pED92 *leu2*:pED94 *WHI5-mCitrine:TRP* ste5Δ::pED147 *ste20Δ*::*NAT* | |

| | | |
|---|---|---|
| yED307 | *HTA2-iRFP-KAN ura3*:pED92 *leu2*:pED94 *WHI5-mCitrine:TRP* ste5Δ::pED147 *ste7Δ*::*NAT* | |
| yED310 | *HTA2-iRFP-KAN ura3*:pED92 *leu2*:pED94 *WHI5-mCitrine:TRP* ste5Δ::pED147 *ste20Δ*::pED159 | S11 |
| yED311 | *HTA2-iRFP-KAN ura3*:pED92 *leu2*:pED94 *WHI5-mCitrine:TRP* ste5Δ::pED147 *ste20Δ*::pED160 | S11 |
| yED312 | *HTA2-iRFP-KAN ura3*:pED92 *leu2*:pED94 *WHI5-mCitrine:TRP* ste5Δ::pED147 *ste20Δ*::pED161 | S11 |
| yED313 | *HTA2-iRFP-KAN ura3*:pED92 *leu2*:pED94 *WHI5-mCitrine:TRP* ste5Δ::pED147 *ste7Δ*::pED162 | S11 |
| yED314 | *HTA2-iRFP-KAN ura3*:pED92 *leu2*:pED94 *WHI5-mCitrine:TRP* ste5Δ::pED147 *ste7Δ*::pED163 | S11 |
| ySP300 | $cdc28_{as}$ | |
| ySP694 | *leu2:*pSP395 | F7 |
| ySP730 | *HTA2-tdiRFP-HIS WHI5-mCitrine:TRP ste5Δ::NAT ura3:pSP365* | |
| ySP737 | *HTA2-tdiRFP-HIS WHI5-mCitrine:TRP ura3:pSP365 ste5Δ::*pED137 | |
| ySP739 | *HTA2-tdiRFP-HIS WHI5-mCitrine:TRP ura3:pSP365 ste5Δ::*pED147 | |
| ySP740 | *HTA2-tdiRFP-HIS WHI5-mCitrine:TRP ura3:pSP365 ste5Δ::*pED137 *bar1Δ*::*NAT* | F6 |
| ySP741 | *HTA2-tdiRFP-HIS WHI5-mCitrine:TRP ura3:pSP365 ste5Δ::*pED138 *bar1Δ*::*NAT* | F6 |
| ySP742 | *HTA2-tdiRFP-HIS WHI5-mCitrine:TRP ura3:pSP365 ste5Δ::*pED147 *bar1Δ*::*NAT* | F6 |
| ySP871 | *HTA2-tdiRFP-LEU WHI5-mCitrine:TRP ura3:pSP365 bar1Δ*::*NAT* | S5, 7 |
| ySP860 | *HTA2-CFP-HIS WHI5-mCitrine:TRP Kss1-mScarlett-LEU bar1-frameshifT30* | S5, 6 |
| ySP865 | *HTA2-CFP-HIS WHI5-mCitrine:TRP Kss1-mScarlett-LEU ste5Δ::*pED137 *bar1-frameshifT30* | F6 |
| ySP866 | *HTA2-CFP-HIS WHI5-mCitrine:TRP Kss1-mScarlett-LEU ste5Δ::*pED147 *bar1-frameshifT30* | F6 |
| ySP867 | *HTA2-CFP-HIS WHI5-mCitrine:TRP Kss1-mScarlett-LEU ste5Δ::*pED138 *bar1-frameshifT30* | F6 |

- The strain ySP694 is derived from the *MAT***α** ancestor ySP122. All remaining strains are descendant of the *MAT***a** ancestor ySP2.

**Supplementary Table1: list of plasmids**

| Plasmid | Insert | Backbone |
|---|---|---|
| pED92 | $P_{RPS6B}$ -$STE7_{1\text{-}33}$ -*NLS-NLS-Cherry* | pSIV URA |
| pED94 | $P_{RPS6B}$ -$STE7_{ND}$ -*NLS-NLS-CFP* | pSIV LEU |
| pED137 | *STE5:HIS* | pGT HIS |
| pED138 | $STE5_{8A}$ *:HIS* | pGT HIS |
| pED141 | $P_{RPS6B}$ -$STE7_{1\text{-}33}$ -*NLS-NLS-mCherry*<br>$P_{RPS6B}$ -$STE7_{ND}$ -*NLS-NLS-CFP* | pSIV URA |
| pED147 | $STE5_{ND}$ *:HIS* | pGT HIS |
| pED159 | $STE20_{ND}$ *:ADE* | pGT ADE |
| pED160 | $STE20_{11A}$ *:ADE* | pGT ADE |
| pED161 | $STE20_{CRIB\Delta}$ *:ADE* | pGT ADE |
| pED162 | $STE7_{ND}$ *:ADE* | pGT ADE |
| pED163 | $STE7_{6A}$ *:ADE* | pGT ADE |
| pED164 | $STE5_{ND}$ *:ADE* | pGT ADE |
| pSP365 | $P_{RPL24A}$-*mCherry-SZ2*<br>$P_{AGA1}$-*NLS-NLS-SZ1* | pSIV URA |
| PSP395 | $P_{RPS2}$-*CFP* | pRS305 |

- NLS represents the peptidic sequence :
  QQMGRGSEFEELGSPLKKLKISPDTASGLV
- *STE7* $_{1\text{-}33}$ represents the peptidic sequence:
  MFQRKTLQRRNLKGLNLNLHPDVGNNGQLQEKT
- *STE7* $_{ND}$ represents the peptidic sequence:
  MFQRKTLQAANLKGANANLHPDVGNNGQLQEKT
- SZ1 represents the peptidic sequence:
  LFFDFLTQIRDFFLQVRDQVFFVQVLFLQGFVLIFQRRNFVFELRNQV
- SZ2 represents the peptidic sequence:
  ARNAYLRKKIARLKKDNLQLERDEQNLEKIIANLRDEIARLENEVASHEQ
- pSIV and pGT backbone from Wosika et al. Molecular Genetics and Genomics, 2016.
- pRS305 backbone from Sikorski et al. Genetics, 1989.
